# Proteomic profiling of baseline CSF and serum from HDClarity identifies signatures for Huntington disease staging and stratification

**DOI:** 10.64898/2026.08.06.743256

**Authors:** Nicholas S. Caron, Inês Caldeira Brás, Jessica C. Barron, Emily M. Harvey, Jeffrey N. Bone, Blair R. Leavitt, Michael R. Hayden

## Abstract

**Background:** Sensitive biomarkers that objectively stage Huntington disease (HD) are needed to improve participant stratification and facilitate the enrichment of clinical trials with biologically and clinically homogeneous populations. The HDClarity study, an international longitudinal biofluid collection initiative for HD, provides a unique resource for large-scale proteomic profiling of matched CSF and serum samples from healthy controls and individuals across premanifest and manifest stages of HD. Here, we leveraged baseline proteomic data from HDClarity to characterize protein signatures associated with disease stage and clinical severity, compare measurements across analytical platforms and biofluid compartments, and identify candidate multi-protein panels for disease staging.

**Methods:** Baseline proteomic data generated using Olink Explore (∼3,000 proteins) and SomaScan (∼7,000 proteins) were analyzed in matched CSF and serum samples from 315 HD gene-expansion carriers and 92 non-HD controls. A total of 2,119 proteins overlapped between Olink and SomaScan, enabling assessment of cross-platform concordance, while CSF-serum relationships were evaluated using all available protein measurements within each assay. Covariate-adjusted linear regression models were used to assess disease stage-associated differences in protein abundance, while partial correlation analyses evaluated relationships between protein abundance, clinical severity in HD gene-expansion carriers, and estimated years to disease onset in premanifest participants. A nested machine-learning pipeline incorporating univariate feature ranking, penalized regression-based feature selection, and repeated cross-validation was used to derive compact multi-protein classifiers for HD staging.

**Results:** Cross-platform and CSF-serum correlations were highly protein-dependent, with some analytes showing strong concordance and others exhibiting weak or inverse relationships. These findings highlight substantial heterogeneity in biomarker behaviour across analytical platforms and biofluids. Adjusted models identified both known HD-associated markers (NEFL, GFAP, CHI3L1) and less well-characterized proteins in CSF (TNFRSF8, TPM3, NPPB) and serum (OMG, NPTXR, NCAN) whose baseline abundance differed across clinical and/or HD-Integrated Staging System (HD-ISS) stages. Partial correlation analyses revealed additional candidate biomarkers associated with clinical severity and years to predicted disease onset. Machine-learning models identified compact CSF and serum protein panels that achieved high classification accuracy (AUC > 0.9) across all clinical contrasts, with models distinguishing the transitions from HD-ISS stage 0 to 1 and from premanifest to early manifest disease demonstrating robust generalization to unseen data.

**Conclusions:** This study provides the first large-scale comparison of matched CSF and serum proteomic profiles from the HDClarity cohort, identifying robust baseline proteomic signatures across the HD continuum. Our findings demonstrate the importance of considering both analytical platform and biofluid when interpreting protein biomarkers and identify compact protein panels with potential utility for objective disease staging, patient stratification, and clinical trial enrichment in HD.

## Background

There is a critical unmet need for sensitive and dynamic biomarkers measurable in accessible biofluids for Huntington disease (HD). Such markers should capture early pathophysiological changes in the brain prior to the onset of overt symptoms, enabling more accurate estimation of disease onset, while also providing objective measures of disease severity that complement clinical assessments. Most importantly, robust biomarkers are needed to stratify participants and predict future disease trajectory in clinical trials.

HD is an autosomal dominant neurodegenerative disorder caused by a CAG trinucleotide repeat expansion in the *HTT* gene (1), resulting in production of a mutant huntingtin protein (HTT) with an expanded polyglutamine tract (2). Although the mutation is present in all cells, HD is characterized by selective neurodegeneration within the brain, primarily affecting neuronal populations in the caudate and putamen, as well as cortical and other subcortical structures (3-8). The mechanisms underlying this selective vulnerability remain incompletely understood but are thought to involve multiple interacting pathogenic processes, including somatic CAG repeat instability (9, 10), transcriptional dysregulation (11, 12), impaired axonal transport and trophic support (13), mitochondrial dysfunction (14), and *HTT1a* transcript production (15, 16), which together contribute to progressive neuronal dysfunction and neurodegeneration (17). Clinically, HD is characterized by a prolonged premanifest phase, with disease onset typically between 40-50 years of age, followed by progressive motor impairment, cognitive deterioration, and psychiatric disturbances (18).

Several protein biomarkers have emerged as promising indicators of disease biology in HD, although only a small number have been extensively studied and validated for use as HD biomarkers. These include neurofilament light chain (NEFL) and mutant HTT. NEFL is an established biomarker of axonal injury in HD, with levels increasing in CSF and blood from premanifest (preHD) to manifest disease and strongly associated with clinical and neuroimaging measures of disease severity (19-26). Mutant HTT, a primary driver of pathology in HD, is elevated in CSF from people with HD and has been used as a pharmacodynamic biomarker of target engagement in trials evaluating HTT lowering therapies (27, 28). Additional proteins, such as PENK (24, 29, 30), PDYN (24, 31), GFAP (32, 33), and CHI3L1 (23, 24, 34) have been reported to change across disease stages. However, validation across independent cohorts and complementary analytical platforms is needed to establish their utility as biomarkers for HD.

HDClarity is an international, multi-site observational biofluid collection initiative built upon the Enroll-HD clinical research platform, a large prospective longitudinal study providing standardized clinical and biological data from individuals with and at risk for HD. Since 2017, HDClarity has collected longitudinal CSF and matched blood samples alongside clinical data from HD gene-expansion carriers (HDGEC), encompassing both preHD and manifest HD, and from non-HD healthy controls (HC) (NCT02855476), providing a well-characterized resource for biomarker discovery. Initial studies have demonstrated the quality and reproducibility of these samples (35) and examined select metabolites from the kynurenine pathway (36); however, large-scale proteomic profiling of HD biofluids has not been reported to date, representing an important gap in biomarker discovery.

In this study, we analyzed baseline CSF and serum proteomic data from 407 HDClarity participants, including 315 HDGEC and 92 HC, generated using two complementary high-throughput platforms, Olink and SomaLogic SomaScan (Soma). Olink uses pairs of oligonucleotide-labeled antibodies that generate DNA reporter molecules following dual target recognition (37), whereas Soma uses chemically modified single-stranded DNA aptamers (SOMAmers) to recognize protein targets (38). Together, these orthogonal affinity-based technologies enable multiplexed quantification of thousands of proteins from small sample volumes while providing complementary coverage of the proteome.

Using matched CSF and serum datasets, we characterized baseline protein abundance across HD stages, assessed associations with clinical measures and estimated years to predicted disease onset (YTO), and evaluated concordance across platforms and biofluid compartments. This cross-sectional design enabled direct comparison of candidate biomarkers across analytical platforms and assessment of central versus peripheral protein signatures.

Finally, we applied a nested cross-validated machine-learning framework combining univariate protein ranking with multivariate feature selection to identify minimal protein panels for classification of HD Integrated Staging System (HD-ISS) stages (39) and composite Unified Huntington’s Disease Rating Scale (cUHDRS)-defined severity groups (40).

## Methods

### Study design

This study was a secondary analysis of HDClarity proteomic (Olink, CHDI Dataset No.: DATA-00000837; DATA-00000842. Soma, CHDI Dataset No.: DATA-00000811) and clinical datasets (PDS3-R2; CHDI Dataset No.: DATA-00001188). The authors did not collect biological samples or generate the proteomic data used in this analysis. Data were made available to the study team for analysis in accordance with applicable data-access and use requirements from the CHDI Foundation, Inc.

Olink and Soma proteomic datasets were derived from 407 baseline CSF–serum sample pairs from participants in the HDClarity study, including 92 HC, 42 early preHD (early preHD), 85 late preHD, 157 early-stage HD (early HD), 20 moderate-stage HD (moderate HD), and 11 advanced-stage HD (advanced HD) participants.

Based on criteria defined in the HDClarity study protocol, HC were participants aged 18-75 years with either no known family history of HD or a family history of HD but a *HTT* CAG < 36. PreHD participants (age 18–75 years) had *HTT* CAG repeat expansions >40 and lacked diagnostic motor features (UHDRS Diagnostic Confidence <4). They were stratified into early preHD (disease burden score [DBS] <250) and late preHD (DBS ≥250). Manifest HD participants (age 21–75 years) had *HTT* CAG repeat expansions >40 and diagnostic motor features (UHDRS Diagnostic Confidence =4), and were categorized by disease stage based on UHDRS Total Functional Capacity (TFC): early HD (TFC 7–13), moderate HD (TFC 3–6), and advanced HD (TFC 0–2). HD-ISS stages were imputed as in (41) based on landmarks defined in (39).

### Protein measurements in biofluids

CSF and serum samples were profiled using two complementary proteomic platforms: Olink Explore 3k (Olink Proteomics) and SomaScan v4.1 (SomaLogic Inc.). Platform-specific calibrators and quality controls were applied according to the manufacturers’ standard workflows to ensure intra- and inter-plate consistency. Olink protein abundance data were provided as normalized protein expression (NPX) values on a log_2_ scale, while Soma data were provided as normalized, log_2_-transformed protein measurements. These log_2_-transformed data were used for all downstream analyses.

The Olink Explore assay measured 2,926 unique protein targets, of which 40 were not detected in the CSF and serum samples. The Soma assay initially included 7,596 aptamer probes. After excluding probes targeting non-human proteins, 7,291 aptamer probes targeting 6,398 unique human proteins remained for analysis. Soma aptamer SpotIDs were annotated with Entrez Gene symbols. For proteins represented by multiple aptamers that were retained in the final analyses, identifiers were disambiguated using numeric suffixes (e.g., PROTEINX, PROTEINX.1, PROTEINX.2).

### Statistical analyses

Statistical analyses were performed in R version 4.5.0 (R Foundation for Statistical Computing, Vienna, Austria) using RStudio. Volcano, balloon, violin, and scatter plots were generated in GraphPad Prism version 11.0.0 using analyzed data exported from R.

### Multiple linear regression analyses

All linear regression analyses were performed using the R *limma* package. Unless otherwise specified, *p*-values were corrected for multiple testing using the Benjamini–Hochberg (BH) method to control the false discovery rate (FDR).

#### Effect of covariates

To assess the influence of biological and demographic variables on protein abundance, multivariate linear regression models were fit for each protein. Normalized protein values were modeled as a function of age (continuous), sex (binary), CAG repeat length (expanded allele), and education level (ISCED). In analyses including all participants, sex and ISCED were treated as categorical variables, and age was modeled continuously. A secondary analysis restricted to HDGEC examined the relationship between CAG repeat length and protein levels independent of diagnostic status, with CAG modeled as a continuous predictor.

For each protein, model coefficients, *p*-values, and R² values were extracted. Multiple testing correction was applied across proteins for each covariate using the BH method (FDR < 0.05). Age and sex showed consistent proteome-wide effects and were therefore retained as covariates in all downstream analyses. ISCED and CAG repeat length demonstrated minimal impact (<10% of proteins) and were excluded from subsequent models.

#### Differential Protein Abundance Analysis

Baseline protein abundance between HDGEC and controls, across clinical stages of HD, and across HD-ISS stages was assessed using the *limma* framework. Protein abundance values were modeled as a function of Group, Sex, and Age using the design formula: ∼ 0 + Group + Sex + Age.

#### HDGEC vs. controls (discovery analysis)

For the discovery comparison, protein-wise linear models were fit using *lmFit()*, followed by empirical Bayes moderation *((eBayes())*, yielding log_2_ fold changes (logFC), moderated t-statistics, and BH-adjusted *p*-values. FDR < 0.1 was used as a significance threshold for this discovery analysis. Proteins were ranked by FDR-value and visualized using a volcano plot. Additionally, the top 25 most differentially abundant proteins were z-score normalized and visualized in a sample-level heatmap generated with *ComplexHeatmap*.

#### Comparisons of baseline protein levels across disease stages

To assess differential abundance across clinical and HD-ISS stages, protein-wise linear models were again fit using *lmFit()*, and all pairwise stage contrasts were generated with *makeContrasts()*. For each contrast, moderated statistics were computed using *eBayes()*, producing covariate-adjusted logFC values that correspond to group differences in a multiple-regression framework. BH-adjusted *p*-values were computed for each contrast, and proteins were ranked according to the minimum FDR observed across all pairwise comparisons. The top 25 proteins showing the most significant differences (FDR < 0.05) were selected for a stage-ordered heatmap using *ComplexHeatmap*. Protein abundance values were adjusted for age and sex using linear regression, and group-preserved residuals for selected CSF and serum proteins were plotted by disease stage.

### Correlation analyses

All correlation analyses were performed using the *ppco*r package, with partial correlations calculated using *pcor.test()*. Unless otherwise specified, models were adjusted for age and sex, and *p*-values were corrected for multiple testing using the BH method (FDR < 0.05).

#### Cross-platform correlations (Olink vs. Soma)

To assess concordance between platforms, partial Pearson correlations were calculated for proteins quantified by both Olink and Soma. For participants with complete metadata, matched protein identifiers were used to align abundance measures across platforms. For each shared protein, partial correlation coefficients (Pearson’s r) and *p*-values were obtained, and FDR correction was applied.

#### CSF–serum correlations

To examine relationships between central and peripheral compartments, partial Pearson correlations were computed for proteins measured in both CSF and serum. Matched CSF–serum pairs from participants with complete metadata were included. For each shared protein, partial correlation coefficients (Pearson’s r) and p-values were obtained, followed by BH correction.

#### Correlations with YTO

To assess associations between protein levels and YTO in preHD participants, partial Pearson correlations were performed with adjustment for sex only. YTO was derived by estimating the predicted age of onset according to (42), using the formula:

Predicted Age at Onset = 21.54 + exp(9.556 – 0.146 × CAG)

Then subtracting age at baseline:

YTO = Predicted age at onset – Age at baseline

For each protein, partial Pearson correlation coefficients (r) and *p*-values were calculated, and BH-adjusted *p*-values were used to determine significance. Proteins were ranked according to the minimum FDR observed across associations with YTO. The top 25 proteins showing the most significant YTO correlations (FDR < 0.05) were z-score normalized and visualized in a sample-level heatmap using *ComplexHeatmap*.

#### Correlations with clinical measures

To evaluate relationships between protein concentrations and clinical outcomes, partial Spearman correlations were performed in HDGEC, adjusting for age and sex. Clinical measures included the Symbol Digit Modalities Test (SDMT), Verbal Fluency Test (VFT), Stroop Color Naming Test (SCNT), Stroop Word Reading Test (SWRT), Total Motor Score (TMS), Total Functional Capacity (TFC), and the composite Unified Huntington’s Disease Rating Scale (cUHDRS). cUHDRS was calculated as described in (40) using:

cUHDRS = [(TFC−10.4)/1.9]−[(TMS−29.7)/14.9]+[(SDMT−28.4)/11.3]+[(SWR−66.1)/20.1]

For each protein–clinical measure pair, the partial Spearman correlation coefficient (ρ), test statistic, and *p*-value were calculated, and BH-adjusted *p*-values were used to determine significance. Proteins were ranked for each clinical measure according to the minimum FDR observed across all pairwise comparisons. The top 25 proteins showing the most significant correlations (FDR < 0.05) were selected for a stage-ordered heatmap using *ComplexHeatmap*. For visualization, cUHDRS and protein abundance values were independently adjusted for age and sex using linear regression, with the mean of the original variable added back to the residuals to preserve the original scale. Adjusted values for proteins showing significant associations with cUHDRS were plotted for selected CSF and serum proteins.

BH-adjusted *p*-values were computed for each contrast, and proteins were ranked according to the minimum FDR observed across all pairwise comparisons. The top 25 proteins showing the most significant differences (FDR < 0.05) were selected for a stage-ordered heatmap using *ComplexHeatmap*.

### Gene ontology enrichment analyses

To identify overrepresented biological pathways from our differentially abundant proteins, we implemented two functional enrichment analyses. First, we used g:Profiler g:GOSt (https://biit.cs.ut.ee/gprofiler/gost) (43, 44) and Enrichr (https://maayanlab.cloud/Enrichr/) (45, 46) to perform non-ranked enrichment analyses of our CSF and serum proteins differentially expressed in HDGEC. For these analyses, all differentially abundant proteins in HDGEC with nominal p values <0.01 were used to determine the top gene ontology (GO) biological process terms from our Olink CSF, Olink serum, Soma CSF, and Soma serum datasets. To increase specificity of the GO Biological Process terms identified for each dataset, term size cut-offs were applied. For Olink CSF, Soma CSF and Soma serum datasets this cut-off was set to 2,500. For Olink serum, no cut-off was set, as no terms less than 2,500 met significance. Next, we performed non-ranked enrichment analyses of our proteins that correlated with predicted YTO of HD. For these analyses, positive and negative correlated proteins with nominal p values <0.001 were used to determine the top GO biological process terms. Term size cut-off was set to 2,500 for all datasets. Functional enrichment results from g:profiler and Enrichr analyses were nearly identical, therefore, results from a single enrichment analysis platform (g:profiler) were used to determine the top 10 GO biological process terms.

### Nested CV machine learning pipeline

All analyses were performed in R using the *glmnet*, *pROC*, *caret*, *tidyverse*, and *doParallel* packages. Parallel computation was used for cross-validation and bootstrap resampling to improve computational efficiency.

#### Overview of analytical framework

All univariate and multivariate analyses were performed within a unified nested cross-validation (CV) framework to ensure unbiased estimation of discriminatory performance and robust feature selection across clinical and HD-ISS stage contrasts.

#### Nested CV design

Model development and evaluation were performed using a five-fold outer CV scheme. For each outer fold, samples were partitioned into training and held-out test sets. All preprocessing, feature ranking, feature selection, and model fitting were performed exclusively within the training data of each outer fold, and predictions were generated only for held-out samples to obtain out-of-fold (OOF) estimates of performance.

Within each outer fold, an inner CV procedure was used for tuning regularisation parameters in penalised regression models.

Age and sex were included as covariates in all multivariate models and were specified as unpenalized terms. These covariates were not considered candidate biomarkers.

#### Univariate protein-level analysis within nested CV

To evaluate single-protein discriminatory performance, univariate logistic regression models were fit for each protein within the training data of each outer fold. For each contrast, univariate models were fit using raw protein abundance values as single predictors, without adjustment for covariates. Non-parametric receiver operating characteristic (ROC) analysis was used to calculate the area under the ROC curve (AUC), and 95% confidence intervals (CI) were derived using the DeLong method (47). Sensitivity and specificity were computed at the optimal threshold defined by the Youden index.

Univariate AUCs were computed independently within each outer fold and aggregated across folds to assess stability and reproducibility of protein-level effects while preserving strict separation between training and test data.

#### Multivariate feature selection and model training

Within each outer training fold, proteins were first ranked using univariate AUC computed on the training subset. The top 100 proteins were retained for downstream modelling to reduce dimensionality and computational burden.

Feature selection was then performed using least absolute shrinkage and selection operator (LASSO)-penalized logistic regression with inner cross-validation used to select the optimal regularization parameter (λ). Proteins with non-zero coefficients at the selected λ were retained as candidate features for that fold. Feature selection was therefore fold-specific, allowing selected protein sets to vary across resamples.

Selection frequency across the five outer folds was used to quantify feature stability and to construct a consensus ranking of proteins.

#### Consensus panel construction and OOF evaluation

For each contrast, candidate biomarker panels were constructed by ordering proteins according to their selection frequency across outer folds. Panel size was increased sequentially from 2 to 20 proteins based on this fixed consensus ranking.

For each panel size, ridge-regularized logistic regression models were independently trained within each outer fold using only features derived from the corresponding training data. Models were then applied to held-out samples to generate OOF predicted probabilities.

Discriminatory performance was assessed using the AUC, computed from pooled OOF predictions across all folds.

#### Performance estimation and uncertainty quantification

Uncertainty in OOF performance was quantified using nonparametric bootstrapping of OOF predicted probabilities (2,000 class-balanced resamples). Bootstrap replicates lacking representation from both outcome classes were excluded from inference. Median bootstrapped AUC values were reported as the primary performance metric.

#### Minimal protein panels and final model fitting

Parsimonious biomarker panels were defined using a tolerance-based criterion relative to the maximum observed OOF performance. The minimal panel was selected as the smallest panel whose OOF AUC was within 0.02 of the best-performing panel size for each contrast. For visualization and reporting of model coefficients, final minimal panels were refit on the full dataset using ridge-regularised logistic regression. ROC curves for minimal panels were derived from these full-data fits.

## Results

### HDClarity study overview and participant baseline characteristics

Matched baseline CSF and serum samples were collected from 315 HDGEC and 92 HC through the HDClarity study (**Fig. 1A**). HDGEC were classified into five clinical groups representing distinct stages along the HD continuum (according to criteria defined in the **Study Design** section), including early preHD (YTO: 20.24 ± 6.48, HD-ISS: 0.41 ± 0.77), late preHD (YTO: 5.73 ± 6.08, HD-ISS: 1.05 ± 1.05), early HD (cUHDRS: 10.66 ± 3.07, HD-ISS: 2.89 ± 0.31), moderate HD (cUHDRS: 3.427 ± 3.24, HD-ISS: 3.00 ± 0), and advanced HD (cUHDRS: -1.492 ± 2.20, HD-ISS: 3.00 ± 0). Baseline demographic and clinical characteristics of HDClarity participants are presented in **Table 1**.

**Fig. 1.**
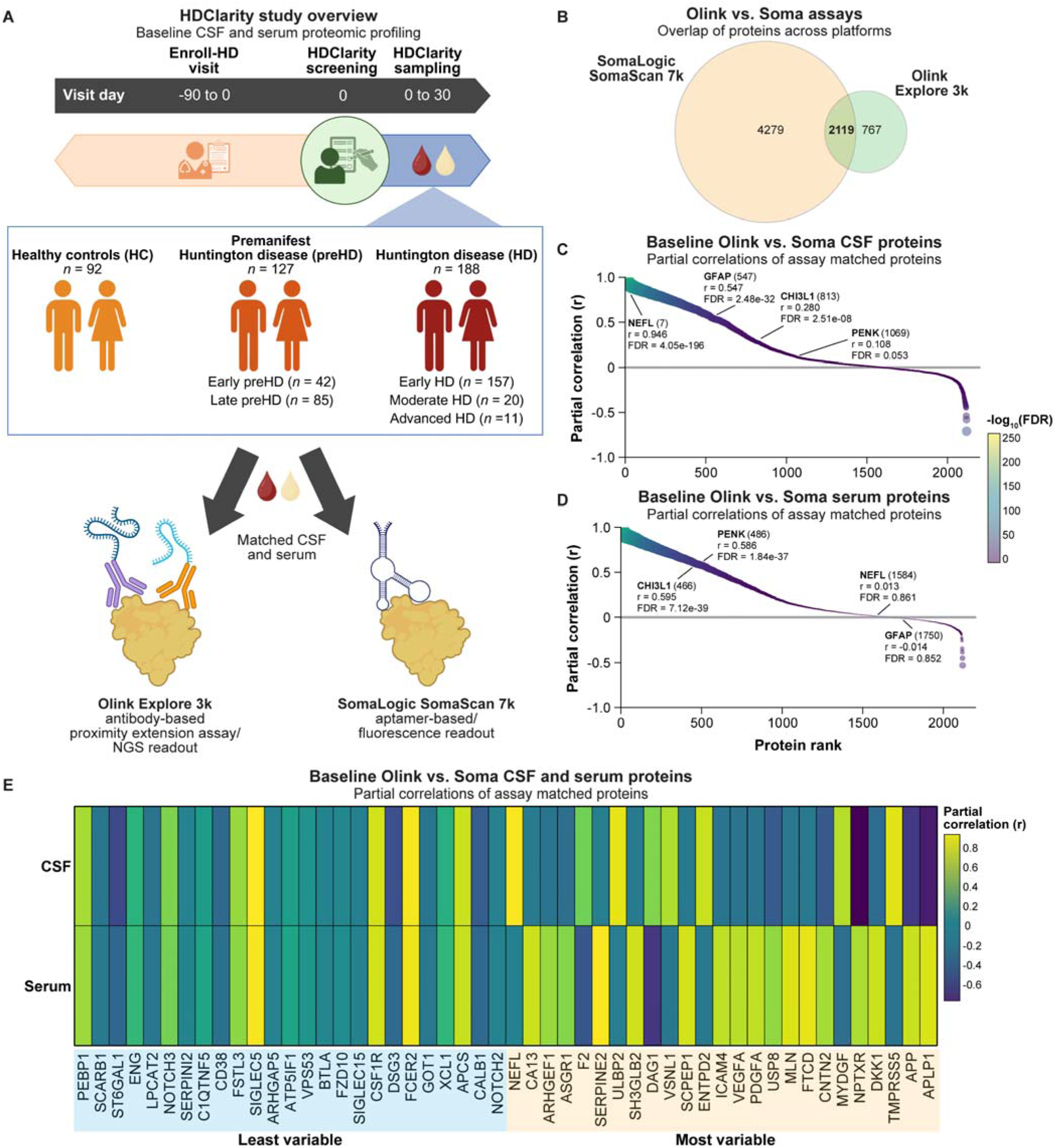
Cross-platform associations of shared CSF and serum proteins (Olink vs Soma). (**A**) Timeline illustrating visit and biofluid sampling contributing to the baseline HDClarity Periodic Dataset 3 (PDS3-R2). All participants in HDClarity were also enrolled in Enroll-HD and attended at least two visits: an HDClarity screening visit and an HDClarity sampling visit, conducted up to 30 days after screening. Matched CSF and serum samples were analyzed from up to 407 participants, including from 92 HC, 127 preHD, and 188 HD participants. Cartoon representation of protein targeting with SomaLogic aptamer-based and Olink proximity extension assays. (**B**) Weighted-Venn diagrams illustrating the number of unique and overlapping proteins measured using Olink and Soma platforms. (**C**-**D**) Balloon plots illustrating partial correlations between assay matched (C) CSF and (D) serum proteins across Olink and Soma platforms, adjusted for age and sex. Proteins were ranked by correlation coefficient. The x-axis shows assay matched protein rank, and the y-axis shows the partial Pearson correlation coefficient (r). Protein rank, correlation coefficient, and FDR-adjusted *p*-value (FDR) for NEFL, GFAP, CHI3L1, and PENK are annotated. (**E**) Heatmap illustrating partial Pearson correlation of the top 25 least and most variable CSF and serum proteins across Olink and Soma platforms.

**Table 1.** HDClarity baseline participant demographics and disease characteristics.

| Characteristic | HC<br>(n = 92) | Early preHD<br>(n = 42) | Late preHD<br>(n = 85) | Early HD<br>(n = 157) | Moderate HD<br>(n = 20) | Advanced HD<br>(n = 11) |
| --- | --- | --- | --- | --- | --- | --- |
| Age (years), mean ± s.d. | 49.28 ± 12.72 | 34.50 ± 7.57 | 41.38 ± 11.35 | 52.27 ± 10.04 | 57.10 ± 9.97 | 56.55 ± 12.20 |
| Sex (M/F) | 36/56 | 17/25 | 42/43 | 89/68 | 12/8 | 5/6 |
| CAG repeat length (high), mean ± s.d. | 19.63 ± 2.99 | 41.55 ± 1.11 | 43.66 ± 2.50 | 43.21 ± 2.48 | 44.00 ± 2.73 | 44.64 ± 4.70 |
| ISCED, mean ± s.d. | 4.12 ± 1.06 | 3.88 ± 1.09 | 3.82 ± 1.09 | 3.88 ± 1.14 | 3.40 ± 1.14 | 3.09 ± 1.14 |
| BMI, mean ± s.d. | 27.96 ± 5.77 (7) | 24.95 ± 5.15 (7) | 24.98 ± 5.27 (15) | 25.06 ± 5.00 (20) | 24.64 ± 7.22 (3) | 24.26 ± 3.90 |
| Years to predicted onset<sup>a</sup>, mean ± s.d. | N/A | 20.23 ± 6.48 | 5.73 ± 6.08 | N/A | N/A | N/A |
| Years since clinical diagnosis, mean ± s.d. | N/A | N/A | N/A | 3.85 ± 3.39 (9) | 8.50 ± 5.19 | 10.45 ± 4.23 |
| HD-ISS<sup>b</sup>, mean ± s.d. | N/A | 0.41 ± 0.77 | 1.05 ± 1.05 (1) | 2.89 ± 0.31 (7) | 3.00 ± 0.00 (1) | 3.00 ± 0.00 (3) |
| DBS<sup>c</sup>, mean ± s.d. | N/A | 204.67 ± 35.47 | 315.40 ± 51.27 | 385.99 ± 79.80 | 463.80 ± 64.35 | 469.82 ± 166.79 |
| CAP100<sup>d</sup>, mean ± s.d. | N/A | 60.71 ± 10.70 | 83.59 ± 13.29 | 103.80 ± 15.02 | 119.81 ± 11.18 | 120.26 ± 21.24 |
| TFC, mean ± s.d. | 12.98 ± 0.15 | 12.76 ± 1.10 | 12.78 ± 0.61 | 10.83 ± 1.63 | 4.85 ± 1.27 | 1.36 ± 0.67 |
| TMS, mean ± s.d. | 1.27 ± 2.15 | 1.33 ± 2.65 | 3.05 ± 3.47 | 27.27 ± 14.15 | 54.42 ± 21.41 (1) | 80.09 ± 18.48 |
| VFT, mean ± s.d. | 23.58 ± 5.20 | 23.38 ± 5.36 | 22.62 ± 5.70 | 15.61 ± 5.79 | 9.30 ± 4.11 | 5.11 ± 2.52 (2) |
| SDMT, mean ± s.d. | 53.27 ± 11.59 | 56.79 ± 9.81 | 52.07 ± 11.82 | 31.50 ± 12.35 | 15.94 ± 11.04 (2) | 6.25 ± 3.10 (7) |
| SWRT, mean ± s.d. | 101.79 ± 15.73 | 100.76 ± 14.55 | 96.66 ± 18.36 | 65.92 ± 19.45 | 37.25 ± 17.99 | 30.88 ± 8.89 (3) |
| SCNT, mean ± s.d. | 78.63 ± 14.81 | 78.26 ± 11.89 | 75.53 ± 13.53 | 51.16 ± 15.57 (1) | 32.15 ± 16.14 | 23.13 ± 8.69 (3) |
| cUHDS<sup>e</sup>, mean ± s.d. | 17.24 ± 1.70 | 17.38 ± 1.56 | 16.65 ± 1.93 | 10.66 ± 3.07 | 3.427 ± 3.24 (3) | -1.492 ± 2.20 (7) |

Continuous variables are presented as mean ± standard deviation (SD). Values shown in red indicate missing data.
<sup>a</sup> YTO = Predicted Age at Onset [= 21.54 + exp(9.556 – 0.146 × CAG)] – Age at baseline
<sup>b</sup> HD-ISS stages were imputed as in (41) based on landmarks defined in (39) .
<sup>c</sup> DBS = Age × (CAG – 35.5)
<sup>d</sup> CAP100 = Age × (CAG – 30)/6.49
<sup>e</sup> cUHDS = [(TFC–10.4)/1.9]–[(TMS–29.7)/14.9]+[(SDMT–28.4)/11.3]+[(SWRT–66.1)/20.1]

The influence of potential biological and demographic covariates on protein levels was assessed using multiple linear regression (**Table S1**). Age, sex, and education level (International Standard Classification of Education; ISCED) were evaluated in all participants, while CAG repeat length was only assessed in HDGEC with CAG >40. Among the 2886 unique human proteins measured using the Olink platform, age was significantly associated with 57.6% of CSF proteins and 42.4% of serum proteins, while sex was associated with 44.7% and 33.0%, respectively (FDR < 0.05). In contrast, CAG repeat length (CSF 2.8%, serum 3.2%) and education (CSF 1.3%, serum 1.0%) were associated with far fewer proteins.

Among the 6,398 unique human proteins measured by Soma (corresponding to 7,291 aptamers), age was associated with 65.6% of CSF and 44.4% of serum proteins, while sex was associated with 39.4% and 19.1%, respectively. Associations with CAG repeat length (CSF 1.1%, serum 0.2%) and education (CSF 1.9%, serum 5.4%) were again limited. Unless otherwise specified, statistical analyses of CSF and serum protein levels measured by both platforms included age and sex as covariates.

### Cross-platform associations of shared CSF and serum proteins (Olink vs Soma)

Of the 6,398 Soma and 2,886 Olink protein targets, 2,119 were shared across platforms (**Fig. 1B**). Concordance between shared proteins was assessed using partial Pearson correlation analysis.

In CSF, 609 proteins showed strong positive correlations (r ≥ 0.5), 302 showed moderate correlations (r = 0.2-0.49), and 1208 showed weak or negative correlations (r < 0.2) (**Fig. 1C**). In serum, 595 proteins showed strong positive correlations, 376 showed moderate correlations, and the remainder showed weak or inverse correlations (**Fig. 1D**).

Examining specific HD-relevant biomarkers between assays, CSF NEFL (r = 0.946, FDR-adjusted *p* = 4.05e-196) and GFAP (r = 0.547, FDR = 2.48e-32) showed a strong positive correlation, CHI3L1 a moderate correlation (r = 0.280, FDR = 2.51e-08), and PENK a weak positive correlation between assays (r = 0.108, FDR = 0.053. In serum, CHI3L1 (r = 0.595, FDR = 7.12e-39) and PENK (r = 0.586, FDR = 1.84e-37) showed strong positive correlations, while NEFL (r = 0.013, FDR = 0.861) and GFAP (r = -0.014, FDR = 0.852) exhibited a weak inverse correlation across platforms. Notably, mutant HTT was not measured by either platform.

Proteins with the greatest and least cross-platform consistency across biofluids were identified by comparing partial correlation coefficients and ranking them by the magnitude of their differences (**Fig. 1E**). Among the least variable proteins – those showing high concordance across both assays and biofluids – PEBP1, SIGLEC5, and FCER2 exhibited consistently strong positive correlations. In contrast, the most variable proteins – those displaying divergent cross-platform performance between CSF and serum – included NEFL, NPTXR, and APP, which showed positive correlations in one biofluid but weak or negative correlations in the other.

### Differential abundance of CSF and serum proteins in HDGEC

Baseline CSF and serum protein levels were compared between HDGEC (including all preHD and HD participants) and HC using multiple linear regression adjusted for age and sex. In CSF, 27 proteins were differentially abundant with Olink (FDR < 0.1), compared with 44 in the Soma dataset (**Fig. 2A**). In serum, 2 proteins were identified with Olink, and 5 with Soma (**Fig. 2B**).

**Fig. 2.**
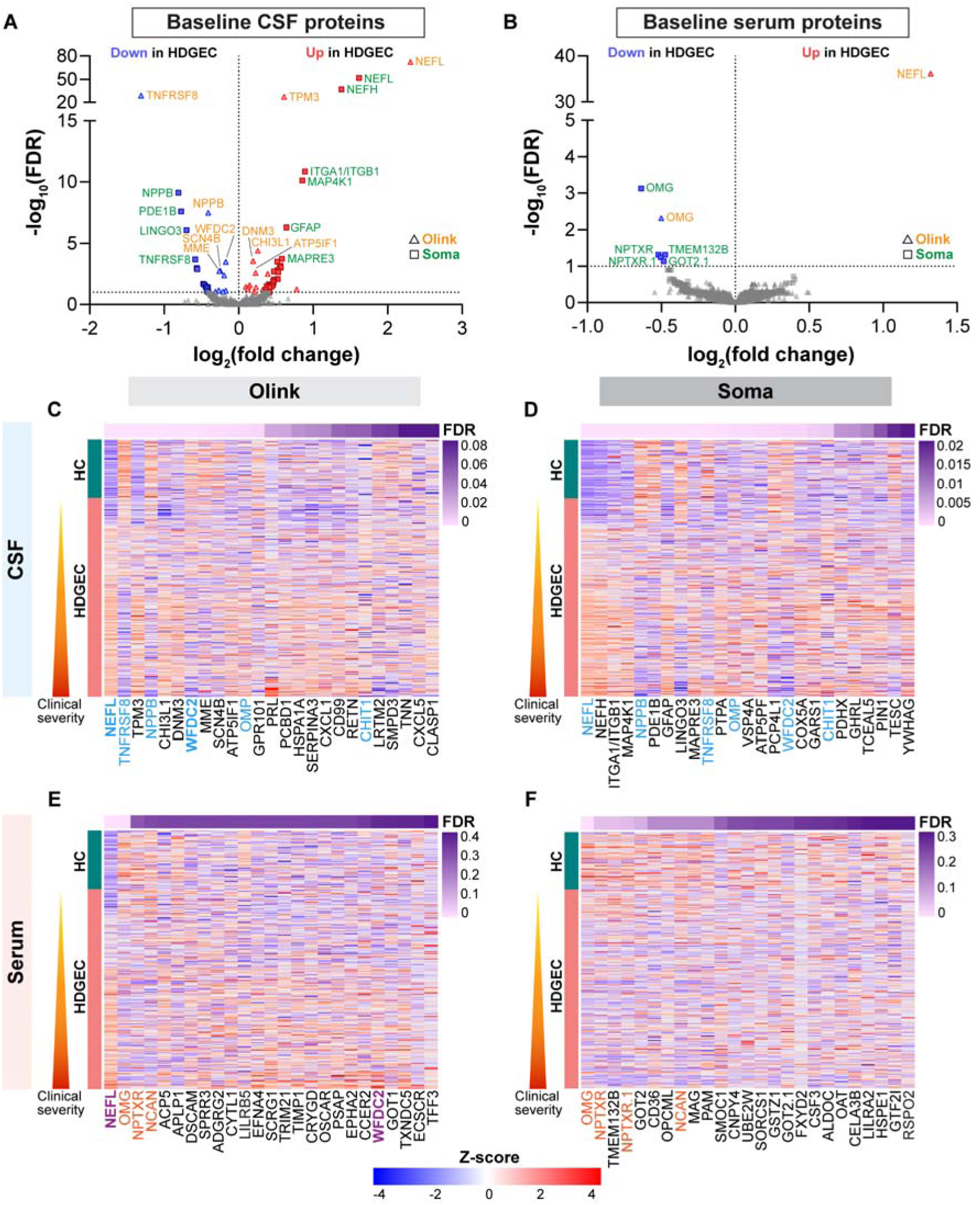
Differential abundance of CSF and serum proteins in HDGEC. (**A**,**B**) Volcano plots showing differential abundance of (A) CSF and (B) serum proteins at baseline between HD gene-expansion carriers (HDGEC) and healthy controls (HC), measured using Olink and Soma platforms, with adjustments for age and sex. Log_2_ fold change is plotted against FDR adjusted *p*-values (FDR; −log_10_ transformed). Olink proteins are depicted as triangles, whereas Soma proteins are depicted as squares. Significantly reduced proteins in each biofluid compared to controls are shown in blue, while those significantly increased are shown in red (FDR < 0.1). The top most significantly reduced and increased Olink (orange) and Soma (green) proteins are annotated. Horizontal dotted line indicates the significance threshold. (**C**-**F**) Sample-level heatmaps of the top 25 differentially abundant proteins at baseline from (C) Olink CSF, (D) Soma CSF, (E) Olink serum, and (F) Soma serum between HDGEC and HC. Linear models were adjusted for age and sex. Columns represent proteins ordered by statistical significance (FDR-adjusted *p*-value), and rows correspond to individual samples grouped by HD mutation status. Cell values correspond to scaled protein abudance levels (z-scores), while annotations display FDR-adjusted *p*-values (FDR) for each protein. Shared CSF proteins between Olink and Soma platforms are shown in blue, and shared serum proteins are shown in red. Shared proteins between biofluid compartments measured on the same platform are shown in bold/purple.

Among the top 25 differentially abundant proteins in CSF, six were shared between the Olink (**Fig. 2C**) and Soma (**Fig. 2D**) platforms – NEFL, TNFRSF8, NPPB, WFDC2, OMP, and CHIT1 – all showing concordant log₂ fold-change (FC) directionality. In the Olink dataset (**Fig. 2C**), NEFL (FC = 2.306, FDR = 4.00e-73), TPM3 (FC = 0.611, FDR = 7.19e-28), and CHI3L1 (FC = 0.258, FDR = 3.90e-05) were the most significantly increased in HDGEC, whereas TNFRSF8 (FC = -1.310, FDR = 1.39e-29), NPPB (FC = -0.408, FDR = 3.00e-08), and WFDC2 (FC = -0.175, FDR = 3.20e-04) were the most significantly decreased.

In the Soma dataset (**Fig. 2D**), NEFL (FC = 1.620, FDR = 1.31e-53), NEFH (FC = 1.381, FDR = 5.93e-38), the ITGA1/ITGB1 complex (FC = 0.893, FDR = 1.05e-11), and MAP4K1 (FC = 0.857, FDR = 6.55e-11) were the most significantly increased relative to HC, while NPPB (FC = -0.810, FDR = 6.65e-10), PDE1B (FC = - 0.771, FDR = 2.42e-08), and LINGO3 (FC = -0.700, FDR = 8.25e-07) were the most significantly reduced.

In serum, only two of the top 25 proteins overlapped between platforms – OMG and NPTXR (detected by two Soma aptamers) – and both showed concordant directionality. In the Olink dataset (**Fig. 2E**), NEFL was the only serum protein significantly elevated in HDGEC (FC = 1.322, FDR = 5.84e-37), while OMG (FC = - 0.500, FDR = 0.005) was the only protein reduced. Notably, NEFL was not altered in the Soma serum dataset (FC = 0.239, FDR = 0.512). In contrast, the top differentially abundant proteins in the Soma dataset, including OMG (FC = -0.637, FDR = 7.21e-04), NPTXR (FC = -0.520, FDR = 0.047), and TMEM132B (FC = -0.474, FDR = 0.047), were all decreased in HDGEC (**Fig. 2F**). Across biofluids, NEFL and WFDC2 were the only differentially abundant proteins among the top Olink-identified candidates in both CSF and serum.

GO enrichment of differentially abundant proteins (nominal p < 0.01) revealed platform-specific biological processes across biofluids. In CSF, Olink proteins were enriched for cytokine-mediated immune signaling, secretion and intracellular transport, and developmental processes, consistent with glial and inflammatory contributions (**Fig. S1A**). In contrast, Soma proteins were enriched for neuronal development, synaptic signaling, and stress responses linked to cell death, suggesting greater sensitivity to neuronal biology (**Fig. S1B**).

In serum, patterns diverged further: Olink proteins were enriched for neurodevelopmental and anatomical structure pathways (**Fig. S1C**), whereas Soma proteins reflected energy and small-molecule metabolism (**Fig. S1D**).

### Differential abundance of CSF and serum proteins across disease stages

Pair-wise comparisons of baseline CSF and serum protein levels across clinical stages were performed using multiple linear regression models adjusted for age and sex. Across clinical disease stage contrasts, ten of the 25 most differentially abundant CSF proteins – NEFL, TNFRSF8, NPPB, PRL, NPTX2, LRTM2, WIF1, CNP, WFDC2, and OMP – were identified across both Olink (**Fig. S2A**) and Soma (**Fig. S2B**) platforms.

In Olink CSF, 123 unique proteins showed one or more significant differences at any disease stage relative to HC (FDR < 0.05). NEFL increased progressively across preHD, with greater abundance in late versus early preHD (**Fig. 3A**; FC = 1.433, FDR = 2.21e-20). TPM3 similarly increased from early to late preHD (**Fig. 3B**; FC = 0.356, FDR = 0.001) and increased further at clinical onset, with greater abundance in early HD than late preHD (FC = 0.260, FDR = 0.001). In contrast, TNFRSF8 (**Fig. 3C**; FC = -1.015, FDR = 1.96e-18), SCN4B (FC = -0.267, FDR = 0.014), MME (FC = -0.275, FDR = 0.025), and NPPB (FC = -0.342, FDR = 6.85e-04) decreased from late preHD to early HD. At later disease stages, PRL increased from moderate to advanced HD (FC = 1.378, FDR = 0.007).

**Fig. 3.**
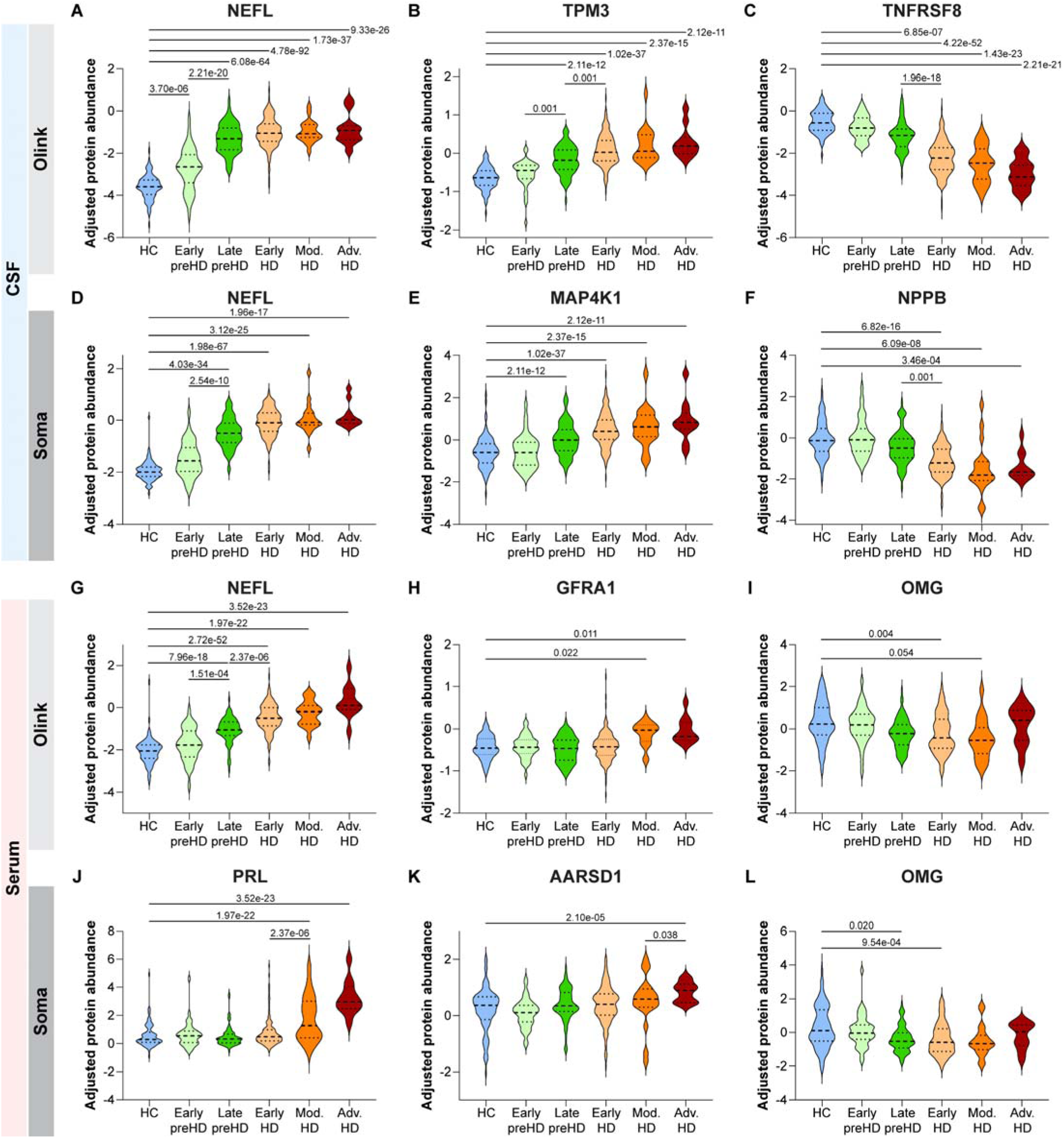
Differential abundance of CSF and serum proteins across clinical stages of HD. Violin plots showing protein abundance across disease stages for (**A-C**) Olink CSF: NEFL, TPM3, and TNFRSF8; (**D-F**) Soma CSF: NEFL, MAP4K1, and NPPB; (**G-I**) Olink serum: NEFL, GFRA1, and OMG; and (**J-L**) Soma serum: PRL, AARSD1, and OMG. Protein abundance values (log_2_-transformed) were adjusted for age and sex while preserving group effects. FDR-corrected *p*-values for significant pairwise contrasts are shown. The dashed line indicates the median, and dotted lines indicate the 25th and 75th percentiles.

The Soma CSF dataset showed a broader set of stage-dependent changes, with more than 500 proteins exhibiting one or more significant differences relative to HC. NEFL (**Fig. 3D**), NEFH, MAP4K1 (**Fig. 3E**), and ITGA1/ITGB1 all increased beginning in late preHD. NEFL (FC = 1.018, FDR = 2.54e-10) and NEFH (FC = 0.888, FDR = 1.46e-06) also increased from early to late preHD, while NPPB declined at the transition from late preHD to early HD (**Fig. 3F**; FC = -0.662, FDR = 0.001). From early to moderate HD, PRL (one of two aptamers; FC = 0.892, FDR = 0.017), CNP (FC = 0.931, FDR = 0.015), and ZFYVE27 increased (FC = 1.416, FDR = 4.88e-05), with PRL increasing further from moderate to advanced HD (one of two aptamers; FC = 1.874, FDR = 1.22e-04).

Stage-dependent changes were also observed in serum, although fewer proteins were affected than in CSF. Among the top 25 differentially abundant serum proteins, PRL and OMG were the only proteins identified across the Olink (**Fig. S2C**) and Soma (**Fig. S2D**) platforms. In the Olink dataset, NEFL, CHGA, and WFDC2 were also differentially abundant in CSF, with CHGA and WFDC2 showing opposite directional changes between compartments. Among 96 unique Olink serum proteins showing one or more stage-dependent differences relative to HC, the majority of changes occurred in advanced HD. NEFL was the only Olink serum protein to show significant differences between consecutive disease stages, increasing from early to late preHD (**Fig. 3G**; FC = 0.680, FDR = 1.51e-04) and again in early HD compared with late preHD (FC = 0.579, FDR = 2.37e-06). PRL (FC = 1.198, FDR = 0.012), GFRA1 (**Fig. 3H**; FC = 0.317, FDR = 0.022), and CCER2 (FC = 0.853, FDR = 0.004) were elevated in moderate HD relative to HC, while OMG was reduced in early HD **(Fig. 3I**; FC = -0.554, FDR = 0.004) and moderate HD compared to HC (FC = - 0.789, FDR = 0.054).

In Soma serum (**Fig. 3J-L**), most significant differences were observed between advanced HD and HC, including AARSD1 (**Fig. 3K**, FC = 1.599, FDR = 2.10e-05) and IL10RB (FC = 1.575, FDR = 1.86e-04). PRL (**Fig. 3J**) and ACP3 were elevated in moderate vs. early HD (PRL: FC = 1.005, FDR = 0.013; ACP3: FC = 1.417, FDR = 1.97e-05), while a broader set of proteins increased from moderate to advanced HD, including PATE1 (FC = 2.073, FDR = 3.65e-04), CARD17 (FC = 1.818, FDR = 4.07e-04), KRAS (FC = 1.824, FDR = 0.005), and PRL (FC = 1.504, FDR = 0.024). In contrast, OMG was decreased in both late preHD (**Fig. 3L**; FC = -0.693, FDR = 0.020) and early HD (FC = -0.698, FDR = 9.54e-04) relative to HC.

Pairwise comparisons of baseline CSF and serum protein abundance across HD-ISS stages (39, 41) were similarly performed using multiple linear regression models adjusted for age and sex. Overall, HD-ISS staging identified many of the same disease-associated protein changes observed across clinical disease stages, particularly in CSF, although fewer proteins showed significant differences between adjacent HD-ISS stages.

Across HD-ISS stage contrasts, eight of the top 25 differentially abundant CSF proteins were detected on both Olink (**Fig. S3A**) and Soma (**Fig. S3B**) platforms. In Olink CSF, NEFL showed progressively greater abundance across HD-ISS stages, increasing from Stage 0 through Stage 3 relative to HC (FC = 1.566, 2.253, 2.082, and 2.580, respectively; **Fig. S3A**). TPM3 similarly showed increased abundance across HD-ISS stages, with significantly greater abundance at Stages 1 (FC = 0.468, FDR = 3.46e-06), 2 (FC = 0.477, FDR = 1.41e-06), and 3 (FC = 0.761, FDR = 2.68e-39) relative to HC. In contrast, TNFRSF8 showed progressively lower abundance beginning at Stage 1 (FC = -0.722, FDR = 6.00e-04), reaching FC = -1.701 at Stage 3 (FDR = 2.01e-52) relative to HC and was also decreased between HD-ISS Stage 2 and Stage 3 (FC = -0.598, FDR = 0.036). Several additional proteins, including NPPB, WFDC2, SCN4B, MME, LRTM2, and WIF1, showed reduced abundance at Stage 3, whereas OMP and PRL were significantly increased.

The Soma CSF dataset showed a similar pattern, with NEFL and NEFH increasing across HD-ISS stages relative to HC and reaching their greatest abundance at Stage 3 (NEFL: FC = 0.897, 1.479, 1.488, and 1.889; NEFH: FC = 0.712, 1.238, 1.198, and 1.636, respectively; **Fig. S3B**). ITGA1/ITGB1 was significantly increased at Stages 2 (FC = 0.973, FDR = 1.89e-04) and 3 (FC = 1.010, FDR = 3.19e-14) compared to HC. NPPB, PDE1B, PCP4L1, LINGO3, WFDC2, TNFRSF8, WIF1, and LRTM2 were reduced at Stage 3 relative to HC, whereas MAP4K1, GFAP, OMP, and PRL were increased at Stage 3. Among adjacent HD-ISS stages, SEMA3E showed a marked reduction at Stage 2 relative to Stage 1, followed by an increase between Stage 2 and Stage 3 (FC = 0.835, FDR = 0.045), further highlighting stage-specific molecular changes within HD-ISS.

In serum, HD-ISS stage-associated differences were less extensive than those observed in CSF. OMG, NCAN, and LILRB5 were among the proteins showing stage-associated changes on both platforms. In the Olink dataset, NEFL showed the most pronounced stage-associated differences, with increased abundance at Stage 0 (FC = 0.584, FDR = 0.002), 1 (FC = 1.032, FDR = 2.07e-10), 2 (FC = 1.010, FDR = 8.41e-10), and 3 (FC = 1.628, FDR = 3.46e-54) relative to HC (**Fig. S3C**). NEFL also increased between HD-ISS Stage 2 and Stage 3 (FC = 0.617, FDR = 0.004). OMG showed reduced abundance at later stages, reaching FC = -0.543 at Stage 3 relative to HC (FDR = 0.004), while LILRB5 was reduced at Stage 2 (FC = -0.745, FDR = 0.090). In the Soma serum dataset, OMG similarly decreased at later HD-ISS stages, reaching FC = -0.688 at Stage 3 relative to HC (**Fig. S3D**; FDR = 0.001). TMEM132B, NCAN, NPTXR (2 aptamers), MAG, and several additional proteins were also reduced at Stage 3, whereas FXYD2 was increased (FC = 0.524, FDR = 0.042). No proteins showed significant differences between adjacent HD-ISS stages.

### CSF-serum protein correlations across analytical platforms

Associations between CSF and serum protein levels were assessed separately for Olink and Soma using partial Pearson correlations adjusted for age and sex. In the Olink dataset, 100 proteins showed strong positive CSF-serum correlations (r > 0.5; **Fig. S4A**), compared with 194 proteins in the Soma dataset (**Fig. S4B**). Among the 25 most strongly correlated proteins, ABO, LILRB5, and LEP were shared across platforms.

For HD-associated proteins differentially abundant in both CSF and serum, NEFL (r = 0.779, FDR = 4.137e-110), REG3A (r = 0.691, FDR = 7.726e-77), and RETN (r = 0.510, FDR = 5.211e-36) demonstrated the highest CSF-serum correlations in the Olink dataset (**Fig. S4C**), whereas REG3A (r = 0.772, FDR = 7.719e-108), **PI3** (r = 0.726, FDR = 7.155e-89), and DEFA5 (r = 0.688, FDR = 2.546e-76) showed the strongest correlations in the Soma dataset (**Fig. S4D**). Of the top 25 differentially abundant proteins with strong CSF-serum correlations, REG3A and PRL were the only proteins shared across platforms.

### Associations of CSF and serum protein abundance with clinical measures of HD

The relationship between baseline biofluid protein levels and clinical measures was assessed in all HDGEC using partial Spearman correlation adjusted for age and sex. Because cUHDRS integrates multiple domains of clinical function, associations with cUHDRS were used as a primary measure of overall clinical status, while associations with individual clinical measures (SCNT, SDMT, SWRT, TFC, TMS, and VFT) were examined to determine whether these relationships extended across clinical domains. Among the top 25 CSF proteins most strongly correlated with clinical outcomes, seven were shared between Olink (**Fig. S5A**) and Soma (**Fig. S5B**).

In the Olink dataset (**Fig. 4A-C**), TNFRSF8, NEFL, NPPB, and TPM3 were significantly associated with all six individual clinical measures and cUHDRS, while WFDC2 was associated with all measures except VFT. TNFRSF8 showed the strongest association with cUHDRS (**Fig. 4A**; ρ = 0.601, FDR = 4.42e-32), followed by NEFL (**Fig. 4B**; ρ = 0.433, FDR = 1.79e-12), with both showing broad associations across individual clinical measures.

**Fig. 4.**
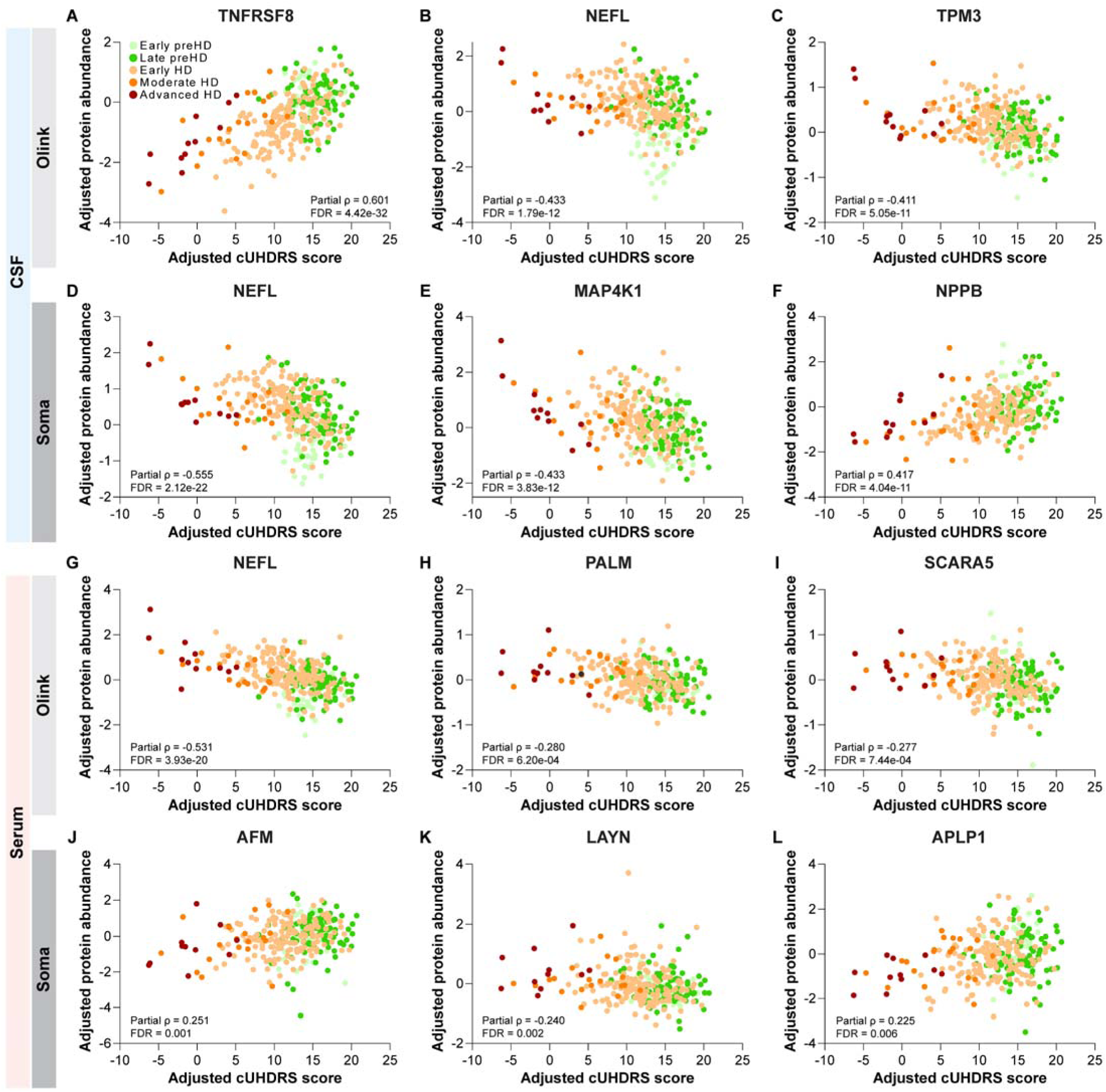
Associations of baseline CSF and serum protein abundance with clinical measures of HD. Scatter plots showing associations between cUHDRS and selected proteins from (**A-C**) Olink CSF: NEFL, TNFRSF8, and TPM3; (**D-F**) Soma CSF: NEFL, MAP4K1, and NPPB; (**G-I**) Olink serum: NEFL, PALM, and SCARA5; and (**J-L**) Soma serum: AFM, LAYN, and APLP1. Protein abundance and cUHDRS values were independently adjusted for age and sex. Points are colored by disease stage. Partial Spearman correlation coefficients (ρ) and FDR-corrected *p*-values are shown for each protein.

In the Soma dataset (**Fig. 4D-F**), NEFL, NEFH, MAP4K1, NPPB, PDE1B, WFDC2, PCP4L1, CNP, and GFAP were significantly associated with all six individual clinical measures and cUHDRS, while WIF1 (two aptamers) was associated with all measures except VFT. NEFL (**Fig. 4D**; ρ = -0.555, FDR = 2.12e-22) and NEFH (ρ = -0.514, FDR = 1.92e-18) showed the strongest associations with cUHDRS. MAP4K1 and NPPB also showed strong associations with cUHDRS (**Fig. 4E,F**; MAP4K1: ρ = -0.433, FDR = 3.83e-12; NPPB: ρ = 0.417, FDR = 4.04e-11).

In serum, fewer proteins were significantly associated with clinical measures, with only C9 and VASN among the top 25 proteins identified by both Olink (**Fig. S5C**) and Soma (**Fig. S5D**). NEFL was the only top Olink protein shared between CSF and serum, with a stronger association with cUHDRS in serum than CSF (serum: ρ = -0.531, FDR = 3.93e-20 vs. CSF: ρ = -0.433). In the Olink serum dataset (**Fig. 4G-I**), NEFL (**Fig. 4G**) and ELN were significantly associated with all six individual clinical measures and cUHDRS, while PALM, ENAH, and PALM2 were associated with all measures except SDMT, and SCARA5 with all measures except VFT. PALM (**Fig. 4H**; ρ = -0.280, FDR = 6.20e-04) and SCARA5 (**Fig. 4I**; ρ = -0.277, FDR = 7.44e-04) also showed significant associations with cUHDRS.

In contrast, Soma serum measurements yielded fewer significant associations overall (**Fig. 4J-L**). AFM (two aptamers) showed the broadest pattern, with significant associations across five of six individual clinical measures and the strongest association with cUHDRS among Soma serum proteins (**Fig. 4J**; ρ = 0.251, FDR = 0.001). LAYN and APLP1 also showed significant associations with cUHDRS (**Fig. 4K,L**; LAYN: ρ = - 0.240, FDR = 2.12e-22; APLP1: ρ = 0.225, FDR = 0.006).

### Associations of baseline CSF and serum protein abundance with YTO

The relationship between baseline biofluid protein levels and YTO in preHD individuals was assessed using partial Pearson correlations adjusted for sex. In CSF, 1073 proteins measured by Olink and 1118 by Soma were significantly associated with YTO (FDR < 0.05; **Fig. S6A**). In serum, far fewer associations were observed, with 46 Olink and 20 Soma proteins reaching significance (**Fig. S6B**).

Among the top 25 CSF proteins most significantly associated with YTO, three – NEFL, CPB1 (detected with two Soma aptamers), and GDF15 – were identified across both Olink (**Fig. S6C**) and Soma (**Fig. S6D**). All top-ranked CSF proteins on both platforms were inversely correlated with YTO, indicating increasing protein abundance as individuals approached predicted disease onset.

In CSF, the strongest Olink associations with YTO included NEFL (r = -0.803, FDR = 3.40e-26), CHI3L1 (r = -0.595, FDR = 2.80e-10), CPB1 (r = -0.581, FDR = 9.13e-10), LTBP2, and TFPI (**Fig. S6C**), while Soma highlighted NEFL (r = -0.777, FDR = 1.28e-22), NEFH (r = -0.726, FDR = 3.79e-18), CBP1 (one of two aptamers; r = -0.567, FDR = 1.33e-08), TRMT6, and CNN1 (**Fig. S6D**).

In serum, IGDCC4 and SCARF2 were shared among the top 25 YTO-associated markers across Olink (**Fig. S6E**) and Soma (**Fig. S6F**), showing consistent correlation directionality between platforms. Several Olink-identified proteins – NEFL, ELN, TFPI, and GDF15 – were significantly correlated with YTO in both CSF and serum and demonstrated concordant directions of association across compartments.

In Olink serum, IGDCC4 (r = 0.443, FDR = 3.06e-04) and COL9A1 were positively correlated with YTO, whereas NEFL (r = -0.665, FDR = 6.11e-15), HAVCR1 (r = 0.409, FDR = 0.001), and HRG showed inverse associations in preHD. In Soma serum, IGDCC4 (r = 0.481, FDR = 1.02e-04), CILP2 (r = 0.444, FDR = 7.53e-04), and MATN4 exhibited the strongest positive correlations, while TAGLN (one of two aptamers; (r = -0.422, FDR = 0.002) and SCARF2 showed the most pronounced negative associations.

GO enrichment of proteins associated with YTO in preHD (nominal *p* < 0.001) revealed shared and platform-specific signatures across biofluids. In CSF, Olink proteins highlighted immune and defense responses, adhesion, migration, and proliferation, reflecting active immune–glial signaling (**Fig. S7A**). Soma CSF proteins similarly pointed to immune and adhesion processes but emphasized immune regulation, wound healing, and motility, indicating coordinated reparative pathways (**Fig. S7B**). In serum, Olink proteins mapped to adhesion, wound healing, and coagulation/hemostasis, consistent with systemic vascular and injury signaling (**Fig. S7C**), while Soma serum proteins focused on protein localization to the plasma membrane, emphasizing cellular trafficking rather than inflammation or coagulation (**Fig. S7D**).

### Classification of HD-ISS stages and cUHDRS-defined severity strata using CSF and serum protein panels

To identify parsimonious proteomic signatures of disease stage, we implemented a nested machine-learning pipeline that combined univariate feature ranking with multivariate feature selection and classification. Within each outer CV fold, protein features were first filtered and ranked according to univariate AUC performance. Top-ranked features were then subjected to LASSO logistic regression for sparse feature selection, and proteins were prioritized based on selection frequency across folds. Ridge-penalized logistic regression models were subsequently trained using incrementally larger protein panels (2–20 proteins), with performance evaluated exclusively on held-out outer folds to generate unbiased OOF AUC estimates. The final consensus biomarker panels were defined using feature stability across folds and refit on the full dataset to demonstrate classification performance.

The highest-ranking individual CSF and serum proteins identified during univariate feature screening are summarized in **Tables S2** and **S3**. Consistent with previous studies, NEFL emerged as one of the strongest individual classifiers across several disease-stage contrasts, particularly in CSF. However, no single protein consistently achieved high classification performance across HD-ISS stage and cUHDRS-defined severity group comparisons, supporting the use of multivariate protein panels.

For HD-ISS stage classification (**Table 2**; **Fig. 5**), proteins most frequently selected across cross-validation (CV) folds are shown for Olink and Soma CSF (**Fig. 5A,D**) and serum (**Fig. 5G,J**). NEFL was among the most consistently selected proteins in CSF across both platforms and contributed to multiple stage contrasts, supporting its robustness as a stage-associated biomarker. TSC22D1, SCP2, CDK2, and ANXA10 also showed high selection stability within individual platforms.

**Fig. 5.**
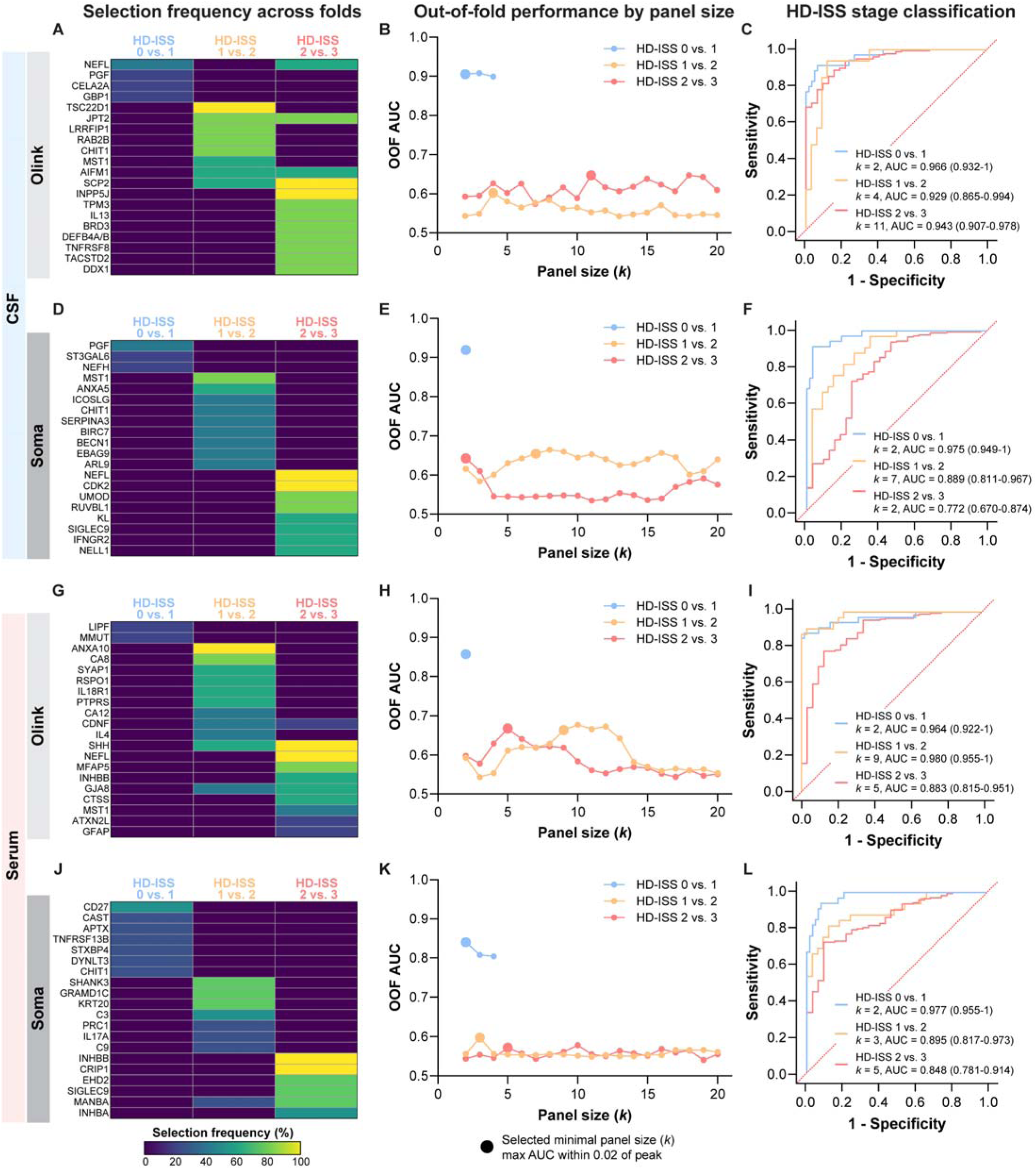
Identification and evaluation of minimal protein panels for classification across HD-ISS stages. (**A-C**) Olink CSF, (**D-F**) Soma CSF, (**G-I**) Olink serum, and (**J-L**) Soma serum. (A, D, G, J) Heatmaps showing the selection frequency of the top 25 proteins for discriminating HD-ISS stage 0 vs. 1 (blue), stage 1 vs. 2 (orange), and stage 2 vs. 3 (red). Proteins were identified using a nested CV framework incorporating univariate feature ranking, LASSO logistic regression, and ridge-penalized classification. Heatmap intensity reflects the percentage of outer CV folds in which each protein was selected, providing a measure of feature selection stability. (B, E, H, K) OOF classification performance as a function of panel size (*k* = 2–20 proteins). Lines represent OOF AUC estimated across five outer CV folds for each HD-ISS stage contrast. Large symbols denote the selected minimal panel size, defined as the smallest panel achieving an OOF AUC within 0.02 of the maximum observed OOF AUC. (C, F, I, L) ROC curves illustrating the classification performance of the selected minimal protein panels refit on the full dataset using ridge-penalized logistic regression. Corresponding panel size (*k*), AUC, and 95% CIs estimated by bootstrap resampling are annotated.

**Table 2.**
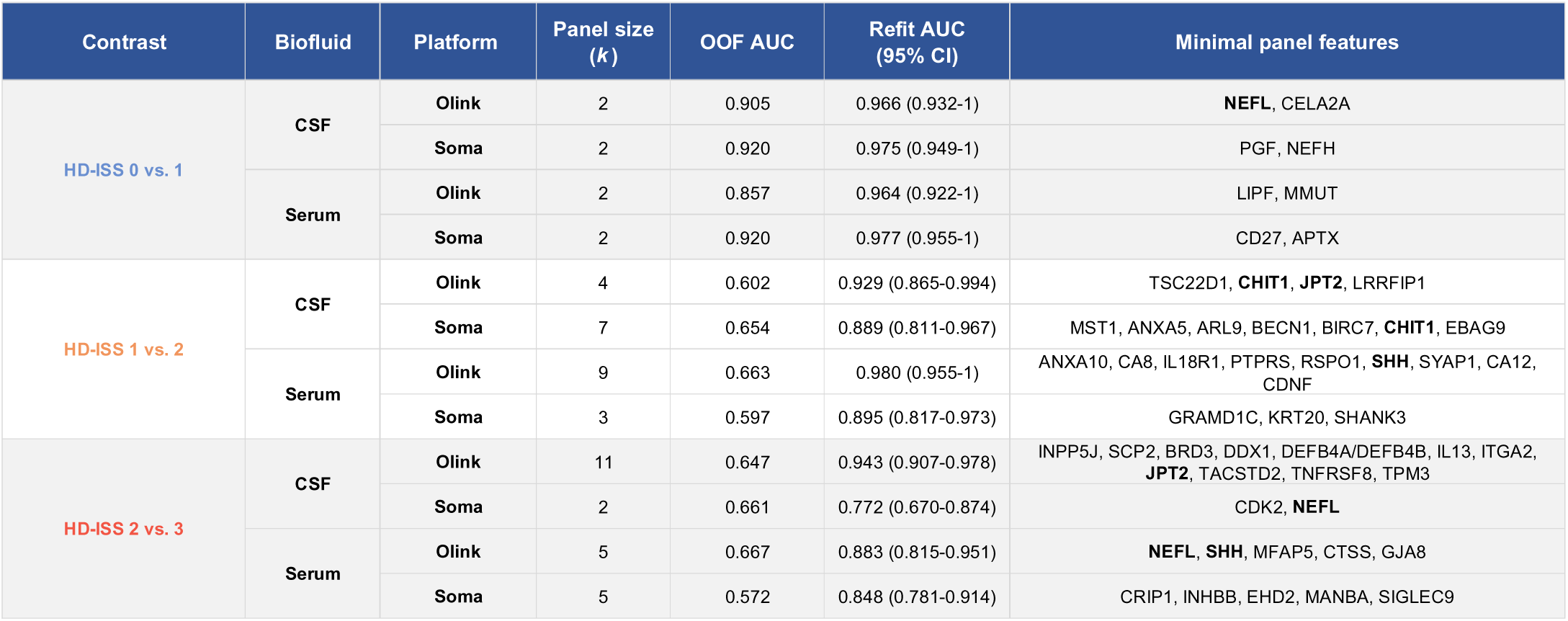
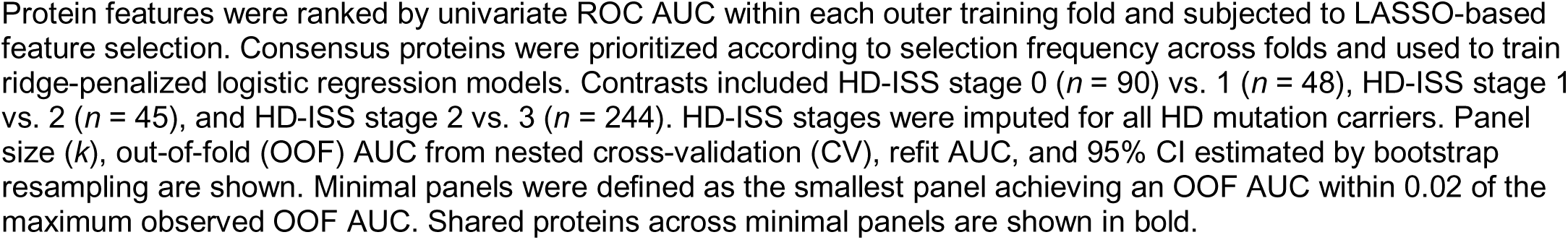
Minimal CSF and serum protein panels for classification across HD-ISS stages.

| Contrast | Biofluid | Platform | Panel size (*k*) | OOF AUC | Refit AUC (95% CI) | Minimal panel features |
| --- | --- | --- | --- | --- | --- | --- |
| HD-ISS 0 vs. 1 | CSF | Olink | 2 | 0.905 | 0.966 (0.932-1) | NEFL, CELA2A |
|  |  | Soma | 2 | 0.920 | 0.975 (0.949-1) | PGF, NEFH |
|  | Serum | Olink | 2 | 0.857 | 0.964 (0.922-1) | LIPF, MMUT |
|  |  | Soma | 2 | 0.920 | 0.977 (0.955-1) | CD27, APTX |
| HD-ISS 1 vs. 2 | CSF | Olink | 4 | 0.602 | 0.929 (0.865-0.994) | TSC22D1, CHIT1, JPT2, LRRFIP1 |
|  |  | Soma | 7 | 0.654 | 0.889 (0.811-0.967) | MST1, ANXA5, ARL9, BECN1, BIRC7, CHIT1, EBAG9 |
|  | Serum | Olink | 9 | 0.663 | 0.980 (0.955-1) | ANXA10, CA8, IL18R1, PTPRS, RSPO1, SHH, SYAP1, CA12, CDNF |
|  |  | Soma | 3 | 0.597 | 0.895 (0.817-0.973) | GRAMD1C, KRT20, SHANK3 |
| HD-ISS 2 vs. 3 | CSF | Olink | 11 | 0.647 | 0.943 (0.907-0.978) | INPP5J, SCP2, BRD3, DDX1, DEFB4A/DEFB4B, IL13, ITGA2, JPT2, TACSTD2, TNFRSF8, TPM3 |
|  |  | Soma | 2 | 0.661 | 0.772 (0.670-0.874) | CDK2, NEFL |
|  | Serum | Olink | 5 | 0.667 | 0.883 (0.815-0.951) | NEFL, SHH, MFAP5, CTSS, GJA8 |
|  |  | Soma | 5 | 0.572 | 0.848 (0.781-0.914) | CRIP1, INHBB, EHD2, MANBA, SIGLEC9 |

Protein features were ranked by univariate ROC AUC within each outer training fold and subjected to LASSO-based feature selection. Consensus proteins were prioritized according to selection frequency across folds and used to train ridge-penalized logistic regression models. Contrasts included HD-ISS stage 0 ( $n = 90$ ) vs. 1 ( $n = 48$ ), HD-ISS stage 1 vs. 2 ( $n = 45$ ), and HD-ISS stage 2 vs. 3 ( $n = 244$ ). HD-ISS stages were imputed for all HD mutation carriers. Panel size ( $k$ ), out-of-fold (OOF) AUC from nested cross-validation (CV), refit AUC, and 95% CI estimated by bootstrap resampling are shown. Minimal panels were defined as the smallest panel achieving an OOF AUC within 0.02 of the maximum observed OOF AUC. Shared proteins across minimal panels are shown in bold.

Evaluation of panel size using OOF performance curves (**Fig. 5B,E,H,K**) identified compact panels with high OOF classification performance. The highest OOF performance was observed for HD-ISS Stage 0 vs. Stage 1, for which two-protein panels achieved AUCs of 0.905 (**Fig. 5B**; Olink) and 0.920 (**Fig. 5E**; Soma) in CSF, and 0.857 (**Fig. 5H**; Olink) and 0.920 (**Fig. 5K**; Soma) in serum. These panels included established HD biomarkers such as NEFL and NEFH, together with CELA2A, PGF, CD27, APTX, LIPF, and MMUT. ROC curves for the selected Stage 0 vs. 1 panels (**Fig. 5C,F,I,L**) showed AUCs ranging from 0.964 to 0.977.

In contrast, classification of HD-ISS Stage 1 vs. Stage 2 and Stage 2 vs. Stage 3 was substantially more challenging, with OOF AUCs of only 0.572–0.667 despite the inclusion of larger protein panels. Although refit AUCs frequently exceeded 0.85, the substantially lower OOF performance indicates limited generalizability of these signatures to unseen data.

To evaluate whether proteomic biomarkers could stratify disease severity independently of formal staging systems, we next examined cUHDRS-defined severity contrasts (**Table 3**; **Fig. 6**). Proteins most frequently selected across cross-validation folds are shown in **Fig. 6A,D** (CSF) and **Fig. 6G,J** (serum). NEFL was recurrently selected across multiple contrasts and platforms, while TNFRSF8, TPM3, GSTT2B, MPP2, and RIPPLY3 also showed high selection stability.

**Fig. 6.**
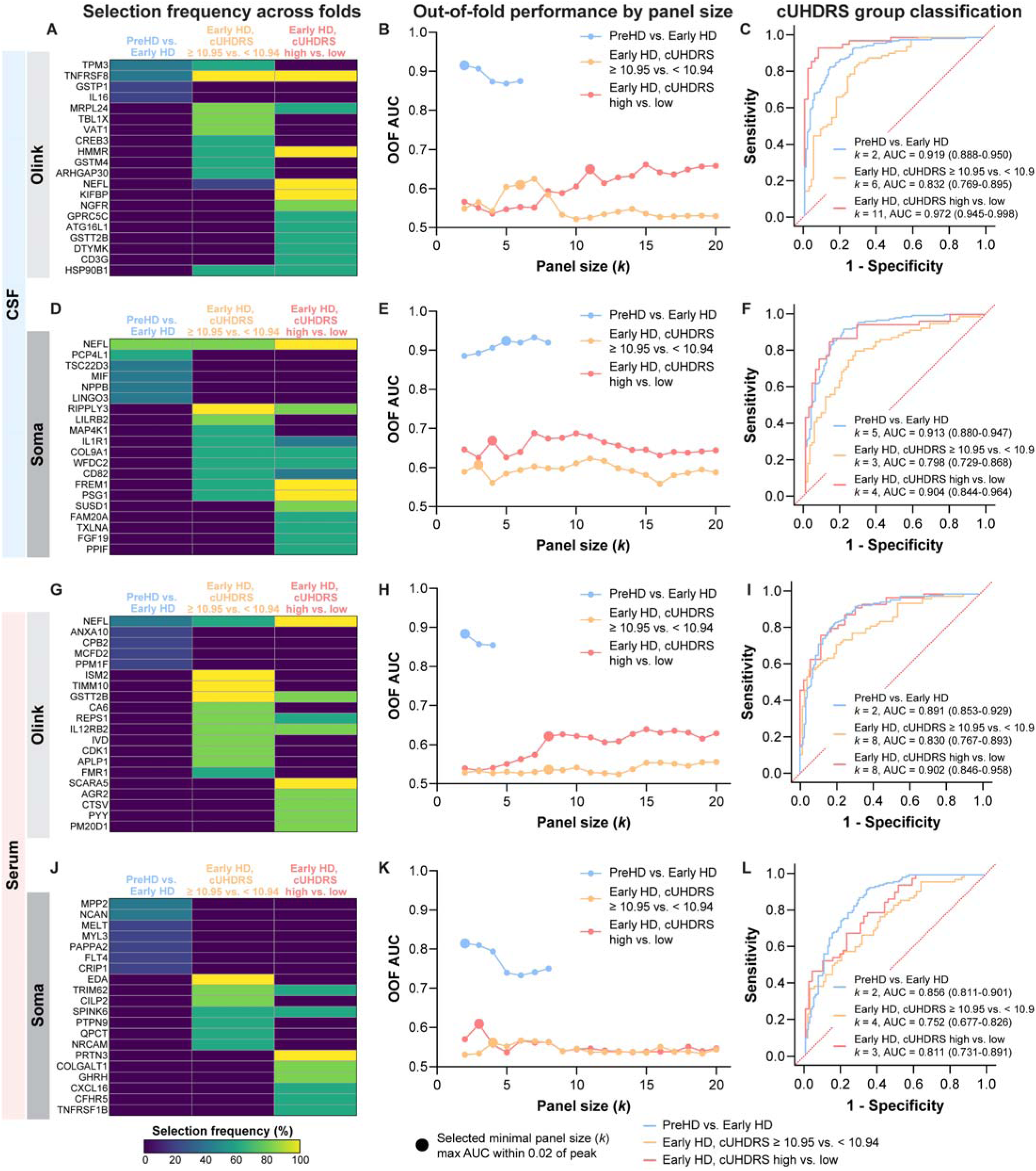
Identification and evaluation of minimal protein panels for classification across cUHDRS-defined severity groups. (**A-C**) Olink CSF, (**D-F**) Soma CSF, (**G-I**) Olink serum, and (**J-L**) Soma serum. (A, D, G, J) Heatmaps showing the selection frequency of the top 25 proteins for discriminating preHD vs. early HD (blue), early HD participants with cUHDRS ≥ 10.95 vs. < 10.94 (orange), and early HD participants in the high vs. low cUHDRS tertile (red). Proteins were identified using a nested CV framework incorporating univariate feature ranking, LASSO logistic regression, and ridge-penalized classification. Heatmap intensity reflects the percentage of outer CV folds in which each protein was selected, providing a measure of feature selection stability. (B, E, H, K) OOF classification performance as a function of panel size (*k* = 2–20 proteins). Lines represent OOF AUC estimated across five outer CV folds for each HD-ISS stage contrast. Large symbols denote the selected minimal panel size, defined as the smallest panel achieving an OOF AUC within 0.02 of the maximum observed OOF AUC. (C, F, I, L) ROC curves illustrating the classification performance of the selected minimal protein panels refit on the full dataset using ridge-penalized logistic regression. Corresponding panel size (*k*), AUC, and 95% CIs estimated by bootstrap resampling are annotated.

**Table 3.** Minimal CSF and serum protein panels for classification across cUHDRS-defined severity groups.

| Contrast | Biofluid | Platform | Panel size ($k$) | OOF AUC | Refit AUC (95% CI) | Panel features |
| --- | --- | --- | --- | --- | --- | --- |
| PreHD vs. Early HD | CSF | Olink | 2 | 0.915 | 0.919 (0.888-0.950) | **TNFRSF8**, TPM3 |
|  |  | Soma | 5 | 0.924 | 0.913 (0.880-0.947) | **NEFL**, PCP4L1, LINGO3, MIF, NPPB |
|  | Serum | Olink | 2 | 0.883 | 0.891 (0.853-0.929) | **NEFL**, ANXA10 |
|  |  | Soma | 2 | 0.819 | 0.856 (0.811-0.901) | MPP2, NCAN |
| Early HD<br>cUHDRS $\geq 10.95$ vs. $< 10.94$ | CSF | Olink | 6 | 0.610 | 0.832 (0.769-0.895) | **TNFRSF8**, MRPL24, TBL1X, VAT1, ARHGAP30, CREB3 |
|  |  | Soma | 3 | 0.607 | 0.798 (0.729-0.868) | RIPPLY3, LILRB2, **NEFL** |
|  | Serum | Olink | 8 | 0.536 | 0.830 (0.767-0.893) | **GSTT2B**, ISM2, TIMM10, APLP1, CA6, CDK1, **IL12RB2**, IVD |
|  |  | Soma | 4 | 0.561 | 0.752 (0.677-0.826) | EDA, CILP2, TRIM62, NRCAM |
| Early HD<br>cUHDRS high vs. low | CSF | Olink | 11 | 0.649 | 0.972 (0.945-0.998) | HMMR, KIFBP, **NEFL**, **TNFRSF8**, NGFR, ATG16L1, CD3G, DTYMK, GPRC5C, **GSTT2B**, HSP90B1 |
|  |  | Soma | 4 | 0.669 | 0.904 (0.844-0.964) | FREM1, **NEFL**, PSG1, RIPPLY3 |
|  | Serum | Olink | 8 | 0.621 | 0.902 (0.846-0.958) | **NEFL**, SCARA5, AGR2, CTSV, **GSTT2B**, **IL12RB2**, PM20D1, PYY |
|  |  | Soma | 3 | 0.609 | 0.811 (0.731-0.891) | PRTN3, COLGALT1, GHRH |

Protein features were ranked by univariate ROC AUC within each outer training fold and subjected to LASSO-based feature selection. Consensus proteins were prioritized according to selection frequency across folds and used to train ridge-penalized logistic regression models. Disease severity contrasts included preHD ( $n = 127$ ) vs. early HD ( $n = 157$ ), early HD participants with cUHDRS scores above ( $n = 79$ ) vs. below ( $n = 78$ ) the cohort median (10.95), and early HD participants in the high ( $n = 52$ ) vs. low cUHDRS tertile ( $n = 52$ ). Panel size ( $k$ ), OOF AUC, refit AUC, and 95% CI estimated by bootstrap resampling are shown. Minimal panels were defined as the smallest panel achieving an OOF AUC within 0.02 of the maximum observed OOF AUC. Shared proteins across minimal panels are shown in bold.

OOF performance curves (**Fig. 6B,E,H,K**) identified compact protein panels for each severity contrast. The strongest classification was observed for preHD vs. early HD, with two- to five-protein panels achieving OOF AUCs of 0.915 (**Fig. 6B**; Olink) and 0.924 (**Fig. 6E**; Soma) in CSF, and 0.883 (**Fig. 6H**; Olink) and 0.819 (**Fig. 6K**; Soma) in serum. ROC curves for these panels ranged from 0.856 to 0.919 (**Fig. 6C,F,I,L**). Notably, Olink achieved this performance using only two proteins in both CSF (**Fig. 6C**; TNFRSF8/TPM3: AUC = 0.919, 95% CI = 0.888-0.950) and serum (**Fig. 6I**; NEFL/ANXA10: AUC = 0.891, 95% CI = 0.853-0.929).

In contrast, classification of early HD participants above vs. below the median cUHDRS score (10.95) showed limited generalizability, with OOF AUCs ranging from 0.536 to 0.610 despite ROC AUCs reaching 0.832. Classification of early HD participants in the highest vs. lowest cUHDRS tertiles showed modestly improved OOF performance, with AUCs ranging from 0.609 to 0.669 across platforms and the highest performance observed in Soma CSF (AUC = 0.669). ROC curves for the selected Olink and Soma CSF panels showed substantially higher AUCs of 0.972 (**Fig. 6C**; 95% CI = 0.945-0.998) and 0.904 (**Fig. 6I**; 95% CI = 0.844-0.964), respectively. The selected panels frequently incorporated NEFL together with proteins involved in immune signaling, cellular stress responses, and neuronal function.

## Discussion

In this study, we present a comprehensive analysis of baseline CSF and serum proteomics from the HDClarity cohort. We identified both known and previously under-explored proteins that were altered across stages of disease and strongly associated with clinical measures and/or YTO in preHD, providing insight into biological processes linked to disease severity and transition to onset. Building on these findings, a nested cross-validated machine-learning framework incorporating univariate protein ranking and multivariate feature selection identified compact protein panels across platforms and biofluids that accurately classified individuals across clinical and HD-ISS stages. These findings highlight the utility of parsimonious protein signatures for objective disease staging. Collectively, these findings support the development of proteomic biomarkers for improved disease staging, patient stratification, and outcome assessment in future HD clinical trials.

NEFL, a well-established marker of axonal injury and the most extensively validated biofluid biomarkers in HD, consistently ranked among the top CSF and serum (Olink) markers across multiple analyses on both Olink and Soma platforms. Interestingly, this pattern was not observed for serum measurements on the Soma platforms, where NEFL failed to show the same disease-related increases, suggesting potential limitations of the Soma assay for measuring serum NEFL, potentially related to assay sensitivity or epitope recognition. NEFL is consistently elevated in CSF and blood, with levels correlating with clinical measures, regional brain atrophy, and is detectable in biofluids decades before predicted onset (19-26, 48). Consistent with these prior studies, NEFL levels in CSF and serum (Olink) were elevated in early preHD (mean YTO ∼20 years), increased progressively with disease severity, and were strongly associated with YTO and clinical measures of HD. NEFL was also recurrently selected in multi-protein panels for classifying across preHD and early HD stages.

NEFH, which was measured only on the Soma platform, showed a remarkably similar pattern across analyses, with disease-stage increases that closely mirrored those of NEFL in CSF. NEFH levels were likewise associated with YTO and multiple clinical measures, supporting its potential as a complementary marker of neuroaxonal injury in HD. These findings reinforce the central role of neurofilament proteins as biomarkers of HD severity and support their utility for disease staging HD and as potential pharmacodynamic markers, particularly given the responsiveness of NEFL to mHTT lowering therapies in preclinical models (49, 50).

Other well-established HD biomarkers, including the glial-enriched inflammatory markers CHI3L1 and GFAP, were elevated in CSF across disease stages and associated with both clinical measures and YTO. These findings are consistent with prior studies (23, 24, 30, 32-34) and further support astroglial activation and inflammatory signaling as early and prominent features of HD pathophysiology.

Additional immune-related proteins, including CHIT1, TNFRSF8, and WFDC2, have also been reported previously and were supported by our findings (29, 34, 51). CHIT1 was increased in CSF across platforms and contributed to panel for distinguishing preHD stages, consistent with early microglia activation in disease. In contrast, TNFRSF8 decreased in CSF with advancing disease stage, showed strong associations with clinical measures, and was a key feature in a panel for classifying late preHD and early HD. WFDC2 was similarly reduced in CSF across disease stages and associated with clinical measures, suggesting altered extracellular protease regulation and possible choroid plexus barrier dysfunction during HD progression.

In contrast to previous reports, PENK and PDYN in CSF did not show disease-associated decreases with either platform (24, 29-31). This discrepancy likely reflects differences in assay design, as mass spectrometry–based studies quantify peptides, whereas Olink and Soma assays measure larger protein regions that may not capture changes in peptide processing or cleavage.

A major contribution of this work is the identification of numerous CSF proteins not previously emphasized in HD biomarker studies. While the breadth of findings precludes detailed discussion of each candidate, several proteins showed consistent and biologically informative associations across platforms, including CNP, CPB1, NPPB, LRTM2, and MMP7. These showed concordant directional changes across datasets, whereas others were platform-specific, highlighting both robust and platform-dependent signals.

A prominent theme that emerged from this analysis was immune and inflammatory dysregulation. Proteins including MIF, IFNG, and MAP4K1 were elevated across disease stages, associated with clinical measures, and contributed to classification models, supporting immune activation in HD pathophysiology.

Extracellular matrix remodeling represented a second theme. Multiple matrix metalloproteinases (MMP7, MMP8, MMP9, MMP12) were elevated across disease stages, while MME was reduced in early HD, suggesting dynamic regulation of protease activity and extracellular matrix turnover.

We also observed consistent alterations in neuronal signaling and integrity. PDE1B, LRTM2, NPPB, NPTX2, and LINGO3 decreased with disease severity, while TPM3 increased and was repeatedly selected in classification models. Notably, OMP showed a strong association with years to onset, supporting early disruption of neuronal function. Together, these findings indicate progressive impairment of neuronal signaling alongside cytoskeletal changes.

Additional alterations were observed in oligodendrocyte and myelin pathways. CNP was increased in moderate vs. early HD and associated with all clinical measures, while JPT2 was elevated in HD-ISS stage 2 vs. stage 1 and contributed to classification panels. CPB1 was associated with YTO in preHD.

Together, these results demonstrate that CSF proteomic changes span immune, extracellular matrix, neuronal, and oligodendrocyte-related pathways, providing a multidimensional view of HD pathobiology.

In serum, fewer proteins showed consistent associations, reflecting biological compartmentalization and reduced CNS signal detection peripherally. Nonetheless, OMG and NCAN decreased with disease severity, consistent with progressive disruption of myelin and extracellular matrix pathways, while NPTXR was reduced, suggesting impaired synaptic maintenance.

In contrast, PRL increased consistently across CSF and serum and platforms. It was also repeatedly selected in multi-protein panels distinguishing disease stages and showed strong cross-compartment concordance, supporting its robustness as a systemic disease marker.

Several additional serum proteins provided complementary information across disease stages. C9 and VASN increased with disease severity and correlated with clinical measures, implicating complement activation and altered tissue remodeling pathways in disease progression. ANXA10 was elevated in early HD and was frequently selected in panels distinguishing preHD from HD and HD-ISS stage 1 from stage 2, while GSTT2B was increased in preHD and contributed to classification of cUHDRS-defined groups in early HD. Together, these proteins suggest that immune, stress-response, and cellular homeostatic pathways are disrupted throughout the disease continuum and may provide useful markers of disease transition and progression.

Together, serum findings identify a smaller but coherent set of proteins reflecting synaptic integrity, extracellular matrix remodeling, immune and stress-response pathways, complementing CSF signals and supporting their utility for disease staging and progression.

Over 1,000 biofluid proteins were significantly associated with YTO among preHD individuals, highlighting extensive molecular changes that precede clinical onset. Rather than acting as deterministic predictors of symptom timing at the individual level, these signals likely reflect progressive, polygenic neurobiological changes that intensify as individuals approach the clinical threshold. As such, proteins associated with predicted onset may provide a quantitative index of proximity to disease manifestation, complementing genetic and clinical estimates. In a clinical research context, these markers could be particularly valuable for identifying individuals closer to onset, enabling enrichment of preventive or early-intervention trials with participants most likely to exhibit measurable change over the study duration. However, prospective validation will be required to determine their stability, specificity, and sensitivity to longitudinal change before consideration for individualized prognostic use.

Notably, YTO associations were far more extensive in CSF than in serum, consistent with CSF more directly reflecting CNS pathology. Among the most robust and reproducible signals across platforms were NEFL, CPB1, and GDF15, all of which increased as individuals approached predicted onset. While NEFL is a well-established marker of neurodegeneration, the consistent identification of CPB1 across platforms, together with its enrichment in oligodendrocytes, implicates early alterations in oligodendrocyte and myelin biology. Other strongly associated CSF proteins, such as CHI3L1 and TFPI, point to immune activation and extracellular matrix or vascular-related processes, suggesting that neuroinflammatory and tissue remodeling pathways are engaged well before clinical onset.

Although fewer proteins were associated with YTO in serum, several showed concordant directionality across compartments, including NEFL, TFPI, and GDF15, supporting their potential as accessible peripheral markers of disease proximity.

Multivariate modeling identified compact CSF and serum protein panels associated with HD stage and disease severity using a nested machine-learning framework that explicitly separated feature selection from performance estimation. This approach provides a more stringent assessment of generalizability by evaluating OOF performance on data not used during model training or feature selection, thereby reducing optimism bias inherent in single-model fitting approaches.

Several important patterns emerged from these analyses. First, the strongest and most reproducible classification performance was observed for transitions corresponding to major biological shifts in disease state, including HD-ISS stage 0 vs. 1 and preHD vs. early HD. These contrasts achieved high OOF AUCs across multiple biofluids and proteomic platforms using remarkably small protein panels, often comprising only two to five proteins. The ability of compact panels to distinguish HD-ISS stage 0 from stage 1 is particularly notable given that this transition is defined by structural neuroimaging evidence of caudate and putamen atrophy. Although striatal volume provides an important measure of disease progression, complementary fluid biomarkers could increase sensitivity to early biological changes and help better align biological disease progression with staging. The strong classification performance observed here supports the presence of measurable proteomic changes in both CSF and serum early in the disease course. Beyond the transition to HD-ISS stage 1, the strong performance observed for preHD vs. early HD suggests that proteomic changes closely track the transition to motor onset, a clinically meaningful milestone for which there are currently no widely accepted fluid-based biomarkers. Together, these findings demonstrate that CSF and serum proteomic profiles capture key biological and clinical disease transitions, including the emergence of measurable striatal atrophy and the onset of motor manifestations.

In contrast, classification of later HD-ISS stage transitions (1 vs. 2, 2 vs. 3) and finer-grained disease severity groups within early HD (cUHDRS ≥ 10.95 vs. < 10.94) proved substantially more challenging. While refit models frequently demonstrated excellent discrimination, OOF performance was more modest, indicating that these transitions are associated with weaker or more heterogeneous proteomic signatures.

This finding is biologically plausible, as later stages of disease likely reflect a convergence of multiple pathological processes and greater inter-individual variability. Importantly, the discrepancy between OOF and refit performance highlights the value of rigorous cross-validation, which can distinguish reproducible signal from model performance that may not generalize to independent datasets.

Despite variability across contrasts, several proteins were repeatedly selected across folds, platforms, and biofluids, including NEFL, as well as proteins linked to immune activation (TNFRSF8, CHIT1), cellular stress (ANXA10, SHH), and neuronal function (NEFH, TPM3). The recurrence of these proteins across independent training iterations supports their robustness as components of disease-associated molecular signatures, while differences in panel composition between contrasts likely reflect the dynamic and stage-specific biology of HD progression. Importantly, even when classification performance was modest in more subtle contrasts, consistent feature selection patterns suggest the presence of underlying biological structure that may become more resolvable with larger cohorts or multimodal integration.

Beyond classification across HD-ISS stages, we evaluated whether proteomic signatures could further refine stratification within clinically relevant groups commonly targeted for clinical trial enrollment. We focused on cUHDRS-defined severity groups because cUHDRS integrates motor, cognitive, and functional measures into a continuous composite outcome and is increasingly used as a sensitive measure of disease progression in HD. Current clinical staging approaches, while widely used to define eligibility and monitor progression, may not fully capture underlying biological heterogeneity among participants with similar levels of clinical impairment. Our findings demonstrate that compact CSF and serum protein panels can distinguish between adjacent severity groups, including transitions relevant to trial inclusion criteria. Proteomic biomarkers could therefore complement clinical measures by capturing molecular differences not reflected in conventional staging systems. Identifying more biologically homogeneous participant subgroups could reduce within-arm variability, improve cohort stratification, and increase statistical power in interventional studies.

Importantly, these proteomic signatures are not intended to replace established clinical frameworks, but rather to augment them by providing an objective molecular layer of disease characterization. Future validation in independent and longitudinal cohorts will be essential to determine whether these signatures improve participant enrichment strategies, track disease progression, or enhance the sensitivity of clinical trials to detect therapeutic effects.

Several limitations of this study should be acknowledged. First, HD-ISS stages were imputed using the approach described in (41), as the imaging data required for direct HD-ISS classification were not available in HDClarity. While this approach enables evaluation of proteomic patterns across biologically defined disease stages, imputed classifications may introduce some uncertainty in stage assignment.

Second, although large-scale proteomic profiling provides important insights into disease-associated biological processes, proteins represent only one component of the broader molecular landscape underlying HD pathogenesis. Integration with complementary modalities, including lipidomics and metabolomics, will be important to capture additional dimensions of cellular dysfunction, such as altered energy metabolism, membrane biology, and signaling pathways that may not be fully reflected at the protein level.

Finally, this study was designed to comprehensively characterize baseline proteomic signatures across analytical platforms, biofluids, and HD disease stages. While these cross-sectional analyses provide insight into the molecular landscape of HD and identify candidate biomarkers associated with disease severity, they do not assess within-person changes over time or establish whether these proteins can reliably track disease progression. Longitudinal analyses will complement these findings by assessing whether these candidates can track disease progression and provide prognostic information in HD.

## Conclusions

Together, these findings establish CSF and serum proteomics as powerful tools for molecular staging and stratification in HD. By integrating cross-platform validation, cross-compartment analyses, and a nested machine-learning framework that separates feature discovery from generalizable performance estimation, this study provides a rigorous framework for biomarker development that balances biological interpretability with translational utility. The consistent identification of compact, reproducible protein signatures across disease stages and severity strata supports the feasibility of proteomic stratification as a complementary approach to clinical assessment. Future work in independent and longitudinal cohorts will be essential to validate these signatures, refine their composition, and determine their utility for tracking disease progression and response to therapeutic intervention. Collectively, these results support the continued development of multi-protein biomarker panels as enabling tools for precision medicine and improved clinical trial design in HD.

## List of abbreviations

AUC: Area under the ROC curve
BH: Benjamini-Hochberg
cUHDRS: Composite Unified Huntington’s Disease Rating Scale
CSF: Cerebrospinal fluid
CV: Cross-validation
FC: Fold change
FDR: False discovery rate
GO: Gene ontology
HC: Healthy controls
HD: Huntington disease
HDGEC: Huntington disease gene-expansion carriers
HD-ISS: HD Integrated Staging System
HTT: Huntingtin
ISCED: International Standard Classification of Education
LASSO: Least absolute shrinkage and selection operator
OOF: Out-of-fold
preHD: Premanifest HD
ROC: Receiver operating characteristic
SCNT: Stroop Color Naming Test
SDMT: Symbol Digit Modalities Test
SWRT: Stroop Word Reading Test
TFC: Total Functional Capacity
TMS: Total Motor Score
VFT: Verbal Fluency Test
YTO: Estimated years to predicted disease onset

## Declarations

### Ethics approval and consent to participate

All analyses were conducted under human research ethics approval from the University of British Columbia (H06-70410) and in accordance with the Tri-Council Policy Statement 2: Course on Research Ethics (TCPS 2: CORE) framework.

### Consent for publication

Not applicable

### Availability of data and materials

HDClarity proteomic (Olink, CHDI Dataset No.: DATA-00000837; DATA-00000842. SomaLogic, CHDI Dataset No.: DATA-00000811) and clinical datasets (PDS3-R2; CHDI Dataset No.: DATA-00001188) underlying this study were provided through Enroll-HD (https://www.enroll-hd.org/for-researchers/) and made available by the CHDI Foundation Inc. and are subject to applicable access and use restrictions. All processed data used to generate figures and tables are available from the corresponding authors on reasonable request.

### Authors’ Contributions

Conceptualization: NSC, MRH

Data analysis: NSC, JNB, ICB, JCB, EH

Visualization: NSC

Writing – original draft: NSC

Writing – review & editing: NSC, ICB, JCB, EH, JNB, BRL, MRH

## Acknowledgements

Data used in this work was generously provided by the participants in the Enroll-HD study and made available by CHDI Foundation, Inc. Enroll-HD is a global clinical research platform intended to accelerate progress towards therapeutics for HD; core datasets are collected annually on all research participants as part of this multi-center longitudinal observational study. Enroll-HD is sponsored by CHDI Foundation, Inc., a non-profit biomedical research organization exclusively dedicated to developing therapeutics for HD. Enroll-HD would not be possible without the vital contribution of the research participants and their families.

HD-Clarity and HD-CSF are biofluid collection initiatives designed to facilitate therapeutic development for HD. HD-Clarity and HD-CSF are led by Dr. Edward Wild and sponsored by University College London. HD-Clarity is funded by CHDI Foundation, Inc., a non-profit biomedical research organization exclusively dedicated to developing therapeutics that will substantially improve the lives of those affected by HD. The Medical Research Council UK (MR/M008592/1) funded HD-CSF. Data used in this work would not be possible without the vital contribution of the research participants and their families in the HD-Clarity and HD-CSF studies.

