## Supplementary materials for "Proteomic profiling of baseline CSF and serum from HDClarity identifies signatures for Huntington disease staging and stratification"

**Table S1. Covariate effects on baseline CSF and serum protein levels across platforms.** Multivariate linear regression models were fitted for each protein to assess associations between normalized protein levels and age, sex, education level (International Standard Classification of Education; ISCED), and CAG repeat length. In analyses including all participants, age was modeled continuously, while sex and ISCED were modeled as categorical variables. A secondary analysis restricted to HD gene expansion carriers (HDGEC) evaluated the association between continuous CAG repeat length and protein levels independent of diagnostic status. The table summarizes the number of proteins significantly associated with each covariate in CSF and serum measured by Olink and SomaScan (Soma) after multiple-testing correction across proteins for each covariate (FDR < 0.05). Columns show the number of proteins (*n*) significantly associated with each covariate (FDR < 0.05) and the corresponding percentage of all proteins analyzed for that platform and biofluid.

| Biofluid | Platform | Covariate | Significant proteins | |
|---|---|---|---|---|
| | | | *n* | % |
| CSF | Olink | Age | 1663 | 57.62 |
| | | Sex (M/F) | 1291 | 44.73 |
| | | CAG (high)* | 82 | 2.84 |
| | | ISCED | 38 | 1.32 |
| | Soma | Age | 4788 | 65.63 |
| | | Sex (M/F) | 2877 | 39.43 |
| | | CAG (high)* | 78 | 1.07 |
| | | ISCED | 140 | 1.92 |
| Serum | Olink | Age | 1223 | 42.38 |
| | | Sex (M/F) | 953 | 33.02 |
| | | CAG (high)* | 93 | 3.22 |
| | | ISCED | 30 | 1.04 |
| | Soma | Age | 3241 | 44.42 |
| | | Sex (M/F) | 1393 | 19.09 |
| | | CAG (high)* | 12 | 0.16 |
| | | ISCED | 392 | 5.37 |

**Table S2. Classification performance of individual CSF and serum proteins across HD-ISS stages.** Top five proteins for each biofluid and analytical platform identified during feature ranking within the nested cross-validation pipeline. Proteins were ranked by univariate ROC AUC within each outer training fold, and the mean rank and median AUC across five outer folds is shown for each HD-ISS stage contrast: HD-ISS stage 0 (*n* = 90) vs. 1 (*n* = 48), HD-ISS stage 1 vs. 2 (*n* = 45), and HD-ISS stage 2 vs. 3 (*n* = 244). HD-ISS stages were imputed for all HD mutation carriers. Proteins shown in bold were identified across multiple stage contrasts, analytical platforms, or biofluids.

| | | 0 vs 1 | | | | | 0+1 vs. 2 | | | | | 0+1 vs 2+3 | | | | |
|---|---|---|---|---|---|---|---|---|---|---|---|---|---|---|---|---|
| **Biofluid** | **Platform** | **HD-ISS stage 0 vs. stage 1** | | | | | **HD-ISS stage 1 vs. stage 2** | | | | | **HD-ISS stage 2 vs. stage 3** | | | | |
| | | **Protein** | **Mean rank (± SD)** | | **Median AUC (± SD)** | | **Protein** | **Mean rank (± SD)** | | **Median AUC (± SD)** | | **Protein** | **Mean rank (± SD)** | | **Median AUC (± SD)** | |
| **CSF** | **Olink** | CCN5 | 2.4 | 1.517 | 0.860 | 0.026 | TSC22D1 | 3.6 | 3.130 | 0.749 | 0.029 | TPM3 | 1.8 | 1.304 | 0.715 | 0.024 |
| | | SOST | 2.6 | 1.517 | 0.864 | 0.011 | JPT2 | 5.0 | 4.637 | 0.742 | 0.032 | SCP2 | 7.6 | 9.236 | 0.691 | 0.018 |
| | | LTBP2 | 4.8 | 1.924 | 0.832 | 0.021 | SCP2 | 8.2 | 5.263 | 0.719 | 0.027 | **NEFL** | 8.0 | 8.246 | 0.685 | 0.017 |
| | | MMP12 | 6.2 | 5.541 | 0.861 | 0.034 | INPP5J | 9.6 | 9.965 | 0.724 | 0.033 | LILRB5 | 21.6 | 20.032 | 0.659 | 0.026 |
| | | **PGF** | 6.2 | 6.301 | 0.837 | 0.022 | THRAP3 | 18.0 | 9.823 | 0.699 | 0.022 | IL13 | 35.0 | 26.125 | 0.647 | 0.034 |
| | **Soma** | SLPI | 4.0 | 3.391 | 0.844 | 0.008 | PRR16 | 11.4 | 17.141 | 0.726 | 0.014 | ANPEP | 8.8 | 5.404 | 0.698 | 0.015 |
| | | CNN1 | 4.6 | 8.050 | 0.856 | 0.027 | ANXA5 | 24.2 | 15.255 | 0.696 | 0.038 | GFAP | 13.8 | 9.257 | 0.692 | 0.015 |
| | | AZGP1 | 8.2 | 4.604 | 0.830 | 0.017 | MYBPC3 | 25.8 | 25.273 | 0.713 | 0.032 | **NEFL** | 14.6 | 18.338 | 0.702 | 0.032 |
| | | **PGF** | 9.4 | 11.589 | 0.841 | 0.017 | MST1 | 34.6 | 43.380 | 0.701 | 0.034 | TRH | 17.4 | 7.232 | 0.682 | 0.024 |
| | | MYL4 | 10.8 | 3.633 | 0.825 | 0.017 | ALPP | 47.6 | 54.326 | 0.698 | 0.020 | NEFH | 18.0 | 14.053 | 0.689 | 0.007 |
| **Serum** | **Olink** | TFPI | 2.6 | 3.050 | 0.797 | 0.031 | ANXA10 | 3.4 | 2.510 | 0.768 | 0.038 | **NEFL** | 1.6 | 1.342 | 0.763 | 0.013 |
| | | **NEFL** | 7.2 | 3.899 | 0.757 | 0.012 | SYAP1 | 8.4 | 4.506 | 0.742 | 0.039 | SHH | 5.2 | 2.864 | 0.705 | 0.027 |
| | | ZP4 | 11.8 | 13.442 | 0.737 | 0.036 | EFNA4 | 17.8 | 11.563 | 0.717 | 0.042 | **INHBB** | 8.4 | 12.095 | 0.708 | 0.017 |
| | | ACRV1 | 15.2 | 15.928 | 0.743 | 0.039 | CKAP4 | 23.8 | 16.053 | 0.699 | 0.058 | PRELP | 11.0 | 10.392 | 0.689 | 0.034 |
| | | LIPF | 19.8 | 25.714 | 0.747 | 0.025 | RSPO1 | 24.0 | 18.588 | 0.717 | 0.035 | ATXN2L | 19.8 | 18.061 | 0.680 | 0.026 |
| | **Soma** | SCUBE1 | 2.0 | 0.000 | 0.804 | 0.016 | PRC1 | 6.0 | 6.285 | 0.763 | 0.020 | **INHBB** | 7.2 | 3.701 | 0.697 | 0.020 |
| | | ARFIP2 | 3.4 | 3.286 | 0.785 | 0.028 | GRAMD1C | 13.4 | 17.300 | 0.768 | 0.030 | EHD2 | 24.6 | 45.676 | 0.703 | 0.021 |
| | | MMP3 | 4.4 | 2.408 | 0.791 | 0.027 | PMP2 | 15.0 | 13.928 | 0.735 | 0.027 | CGB3 | 25.4 | 21.916 | 0.685 | 0.009 |
| | | SFRP4 | 5.4 | 3.209 | 0.773 | 0.005 | SHANK3 | 16.8 | 13.846 | 0.731 | 0.018 | HYOU1 | 39.0 | 57.446 | 0.687 | 0.018 |
| | | KIAA0040 | 7.0 | 5.244 | 0.756 | 0.029 | ENPP5 | 27.4 | 32.020 | 0.727 | 0.055 | CCL7 | 42.8 | 55.549 | 0.687 | 0.026 |

**Table S3. Classification performance of individual CSF and serum protein across disease severity strata.** Top five proteins for each biofluid and analytical platform identified during feature ranking within the nested cross-validation pipeline. Proteins were ranked by univariate ROC AUC within each outer training fold, and the mean rank and median AUC across five outer folds is shown for each disease severity contrast: preHD (*n* = 127) vs. early HD (*n* = 157), early HD participants with cUHDRS scores above (*n* = 79) vs. below (*n* = 78) the cohort median (10.95), and early HD participants in the highest (*n* = 52) vs. lowest cUHDRS tertile (*n* = 52). Proteins shown in bold were identified across multiple disease severity contrasts, analytical platforms, or biofluids.

| Biofluid | Platform | PreHD vs. Early HD | | | | | Early HD cUHDRS ≥ 10.95 vs. < 10.94 | | | | | Early HD cUHDRS high vs. low | | | | |
|---|---|---|---|---|---|---|---|---|---|---|---|---|---|---|---|---|
| | | Protein | Mean rank (± SD) | | Median AUC (± SD) | | Protein | Mean rank (± SD) | | Median AUC (± SD) | | Protein | Mean rank (± SD) | | Median AUC (± SD) | |
| CSF | Olink | **NEFL** | 1.0 | 0.000 | 0.810 | 0.008 | **TNFRSF8** | 1.0 | 0.000 | 0.712 | 0.029 | **TNFRSF8** | 2.0 | 1.732 | 0.724 | 0.029 |
| | | TPM3 | 2.2 | 0.447 | 0.781 | 0.012 | **NEFL** | 9.8 | 7.887 | 0.652 | 0.029 | KIFBP | 7.6 | 6.269 | 0.678 | 0.019 |
| | | SDC4 | 4.0 | 0.707 | 0.743 | 0.012 | MRPL24 | 15.8 | 12.377 | 0.638 | 0.019 | KIAA1549L | 10.8 | 8.643 | 0.673 | 0.023 |
| | | CHI3L1 | 4.4 | 1.140 | 0.751 | 0.014 | MYBPC1 | 29.2 | 26.129 | 0.633 | 0.019 | HMMR | 20.6 | 11.393 | 0.656 | 0.015 |
| | | **TNFRSF8** | 4.6 | 2.702 | 0.772 | 0.023 | TBL1X | 35.2 | 14.342 | 0.623 | 0.011 | GPRC5C | 24.2 | 16.604 | 0.661 | 0.027 |
| | Soma | **NEFL** | 1.0 | 0.000 | 0.832 | 0.012 | **NEFL** | 1.0 | 0.000 | 0.685 | 0.025 | **NEFL** | 1.4 | 0.894 | 0.722 | 0.017 |
| | | **NEFH** | 2.0 | 0.000 | 0.800 | 0.014 | **NEFH** | 6.6 | 6.025 | 0.641 | 0.016 | **NEFH** | 6.2 | 4.712 | 0.702 | 0.017 |
| | | GPT | 5.0 | 2.550 | 0.746 | 0.025 | RIPPLY3 | 33.4 | 29.855 | 0.627 | 0.021 | FREM1 | 9.6 | 11.104 | 0.696 | 0.019 |
| | | GFAP | 5.8 | 5.718 | 0.761 | 0.010 | MAP4K1 | 33.8 | 41.889 | 0.623 | 0.030 | MAP4K1 | 18.6 | 14.571 | 0.669 | 0.021 |
| | | SDHAF2 | 10.0 | 7.106 | 0.735 | 0.026 | IL1R1 | 61.6 | 98.093 | 0.617 | 0.030 | WFDC2 | 29.6 | 15.518 | 0.665 | 0.014 |
| Serum | Olink | **NEFL** | 1.0 | 0.000 | 0.859 | 0.011 | ISM2 | 4.4 | 3.975 | 0.656 | 0.033 | AGR2 | 8.8 | 3.421 | 0.681 | 0.008 |
| | | ELN | 2.0 | 0.000 | 0.780 | 0.007 | TIMM10 | 6.0 | 4.637 | 0.661 | 0.013 | **NEFL** | 12.4 | 10.090 | 0.673 | 0.017 |
| | | LMOD1 | 4.6 | 1.517 | 0.748 | 0.005 | **GSTT2B** | 16.0 | 8.689 | 0.638 | 0.012 | CTSV | 12.6 | 15.662 | 0.678 | 0.033 |
| | | EDA2R | 6.4 | 3.362 | 0.745 | 0.014 | ANXA4 | 18.4 | 11.149 | 0.632 | 0.022 | **GSTT2B** | 13.0 | 20.359 | 0.690 | 0.025 |
| | | CXCL17 | 6.8 | 5.762 | 0.750 | 0.014 | FGF12 | 19.8 | 8.701 | 0.636 | 0.012 | SCARA5 | 13.2 | 7.918 | 0.671 | 0.013 |
| | Soma | PDLIM3 | 2.2 | 2.683 | 0.733 | 0.019 | **MFAP2** | 14.600 | 16.041 | 0.659 | 0.019 | PRTN3 | 23.0 | 16.171 | 0.675 | 0.029 |
| | | TAGLN | 2.6 | 1.342 | 0.726 | 0.016 | **SPINK6** | 32.400 | 29.830 | 0.653 | 0.012 | **SPINK6** | 32.8 | 31.236 | 0.681 | 0.019 |
| | | ASB9 | 4.8 | 3.271 | 0.724 | 0.025 | NARF | 39.600 | 32.176 | 0.633 | 0.015 | CXCL16 | 44.6 | 44.506 | 0.683 | 0.017 |
| | | EHD2 | 8.6 | 4.615 | 0.702 | 0.011 | EDA | 39.800 | 40.929 | 0.651 | 0.023 | COLGALT1 | 46.2 | 69.525 | 0.704 | 0.037 |
| | | TPPP3 | 8.6 | 7.893 | 0.699 | 0.024 | TRIM62 | 43.800 | 83.443 | 0.673 | 0.035 | **MFAP2** | 48.0 | 80.281 | 0.685 | 0.028 |

**Fig. S1. Functional enrichment analysis of differentially abundant CSF and serum proteins in HDGEC.** (**A-D**) Balloon plots illustrating enrichment of the top 10 Gene Ontology (GO) Biological Process terms among proteins differentially abundant in HDGEC compared with HC for: (A) Olink CSF, (B) Soma CSF, (C) Olink serum, and (D) Soma serum. The y-axis shows the top 10 enriched Biological Process terms, and x-axis shows -log10 FDR-adjusted *p*-value. Enrichment was performed using g:GOSt. Proteins with nominal *p*-values <0.01 from the differential abundance analysis (**Fig. 2**) were used as input. The term size cutoff was set to 2,500, except for Olink serum, where only terms >2,500 met significance. Balloon colour indicates term size, and balloon size reflects intersection size.

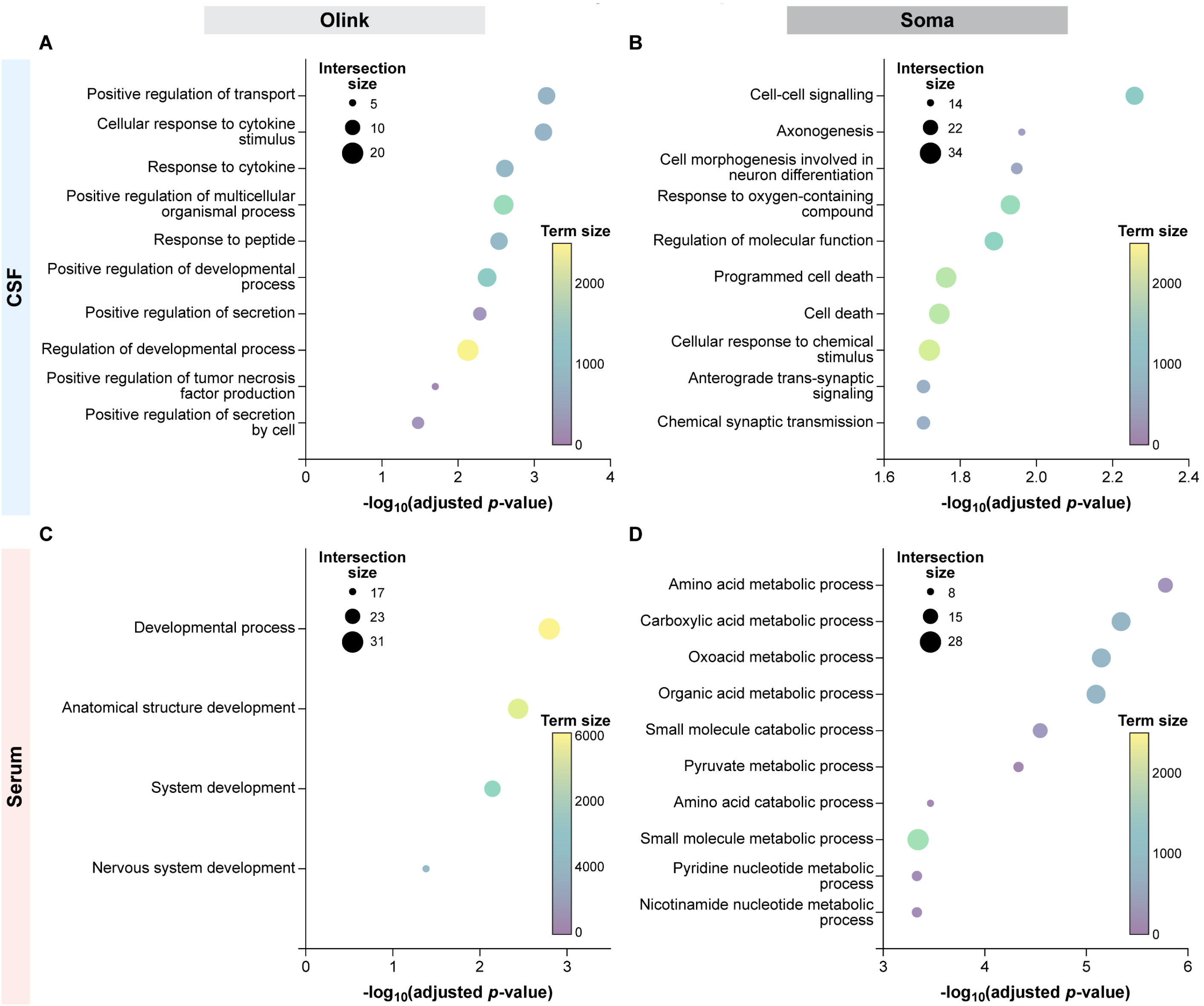

**Fig. S2. Differential abundance of CSF and serum proteins across clinical stages of HD.** (**A-D**) Heatmaps showing the top 25 differentially abundant proteins from (A) Olink CSF, (B) Soma CSF, (C) Olink serum, and (D) Soma serum across pairwise clinical HD stage contrasts. Multiple linear regression models were adjusted for age and sex. Columns represent group contrasts, and rows show proteins ordered first by the number of significant contrasts (FDR-adjusted *p* < 0.05) and then by minimum adjusted *p*-value. Cell values correspond to $\log_2$ fold changes, with text indicating the direction and magnitude of change, while colour intensity reflects statistical significance (FDR-adjusted *p*-value). Shared CSF proteins between Olink and Soma platforms are shown in blue, and shared serum proteins are shown in red. Shared proteins between biofluid compartments measured on the same platform are shown in bold/purple. HC: healthy controls

**Olink** | **Soma**

**A** (CSF, Olink)

| | vs. HC: Early preHD | vs. HC: Late preHD | vs. HC: Early HD | vs. HC: Mod. HD | vs. HC: Adv. HD | Late/ Early preHD | Early HD/ Late preHD | Mod./ Early HD | Adv./ Mod. HD |
|---|---|---|---|---|---|---|---|---|---|
| NEFL | 0.86 | 2.29 | 2.58 | 2.63 | 2.72 | 1.43 | 0.28 | 0.05 | 0.10 |
| TNFRSF8 | −0.26 | −0.68 | −1.69 | −1.99 | −2.43 | −0.42 | −1.02 | −0.30 | −0.43 |
| TPM3 | 0.12 | 0.48 | 0.74 | 0.85 | 0.93 | 0.36 | 0.26 | 0.11 | 0.09 |
| SCN4B | −0.03 | −0.07 | −0.33 | −0.42 | −0.64 | −0.04 | −0.27 | −0.09 | −0.22 |
| MME | −0.06 | −0.09 | −0.37 | −0.38 | −0.34 | −0.04 | −0.28 | −0.01 | 0.04 |
| NPPB | 0.02 | −0.18 | −0.52 | −0.83 | −0.95 | −0.20 | −0.34 | −0.30 | −0.13 |
| **PRL** | 0.12 | 0.19 | 0.33 | 1.03 | 2.40 | 0.07 | 0.14 | 0.70 | 1.38 |
| NPTX2 | −0.03 | −0.04 | −0.19 | −0.51 | −0.91 | −0.02 | −0.14 | −0.32 | −0.40 |
| CHI3L1 | 0.02 | 0.28 | 0.29 | 0.44 | 0.09 | 0.26 | 0.01 | 0.15 | −0.35 |
| LRTM2 | −0.07 | −0.05 | −0.29 | −0.71 | −1.18 | 0.02 | −0.24 | −0.41 | −0.47 |
| WIF1 | −0.03 | −0.11 | −0.33 | −0.78 | −1.25 | −0.08 | −0.23 | −0.45 | −0.47 |
| ADAM23 | −0.02 | 0.03 | −0.09 | −0.26 | −0.42 | 0.05 | −0.12 | −0.16 | −0.17 |
| CNP | −0.07 | 0.09 | 0.11 | 0.46 | 0.82 | 0.16 | 0.02 | 0.35 | 0.36 |
| **WFDC2** | −0.07 | −0.06 | −0.20 | −0.32 | −0.57 | 0.01 | −0.14 | −0.12 | −0.25 |
| DNM3 | 0.01 | 0.13 | 0.25 | 0.27 | 0.23 | 0.12 | 0.12 | 0.02 | −0.04 |
| PAM | −0.03 | 0.07 | −0.11 | −0.35 | −0.74 | 0.10 | −0.19 | −0.24 | −0.39 |
| CBLN4 | −0.07 | 0.00 | −0.07 | −0.46 | −0.88 | 0.07 | −0.08 | −0.38 | −0.42 |
| GPR158 | −0.02 | −0.00 | −0.03 | −0.08 | −0.25 | 0.01 | −0.03 | −0.05 | −0.16 |
| MANEAL | 0.04 | 0.11 | −0.08 | −0.25 | −0.85 | 0.07 | −0.18 | −0.17 | −0.60 |
| ATP5IF1 | 0.06 | 0.20 | 0.26 | 0.29 | 0.34 | 0.13 | 0.07 | 0.02 | 0.06 |
| VGF | −0.04 | 0.05 | −0.05 | −0.28 | −0.69 | 0.09 | −0.10 | −0.23 | −0.41 |
| CSPG5 | −0.05 | 0.05 | −0.05 | −0.23 | −0.48 | 0.10 | −0.10 | −0.19 | −0.25 |
| OMP | 0.23 | 0.25 | 0.47 | 0.46 | 0.57 | 0.02 | 0.22 | −0.01 | 0.11 |
| NPY | 0.04 | 0.08 | 0.01 | −0.22 | −0.66 | 0.03 | −0.07 | −0.24 | −0.43 |
| CHGA | −0.00 | 0.03 | −0.04 | −0.09 | −0.24 | 0.03 | −0.07 | −0.05 | −0.15 |

**B** (CSF, Soma)

| | vs. HC: Early preHD | vs. HC: Late preHD | vs. HC: Early HD | vs. HC: Mod. HD | vs. HC: Adv. HD | Late/ Early preHD | Early HD/ Late preHD | Mod./ Early HD | Adv./ Mod. HD |
|---|---|---|---|---|---|---|---|---|---|
| NEFL | 0.45 | 1.47 | 1.88 | 2.01 | 2.12 | 1.02 | 0.41 | 0.14 | 0.11 |
| NEFH | 0.32 | 1.20 | 1.58 | 2.06 | 1.99 | 0.89 | 0.38 | 0.48 | −0.07 |
| **PRL** | 0.07 | 0.13 | 0.37 | 1.27 | 3.14 | 0.06 | 0.25 | 0.89 | 1.87 |
| **PRL.1** | 0.06 | 0.09 | 0.31 | 1.17 | 2.97 | 0.03 | 0.21 | 0.86 | 1.80 |
| NPPB | 0.03 | −0.41 | −1.07 | −1.48 | −1.36 | −0.45 | −0.66 | −0.40 | 0.12 |
| MAP4K1 | 0.01 | 0.69 | 1.05 | 1.18 | 1.46 | 0.69 | 0.35 | 0.13 | 0.28 |
| ITGA1/ITGB1 | 0.39 | 0.73 | 1.03 | 1.07 | 1.38 | 0.34 | 0.30 | 0.04 | 0.31 |
| PDE1B | −0.12 | −0.42 | −0.99 | −1.02 | −1.70 | −0.30 | −0.57 | −0.03 | −0.68 |
| GFAP | 0.10 | 0.45 | 0.76 | 1.04 | 1.27 | 0.35 | 0.31 | 0.28 | 0.23 |
| LINGO3 | −0.15 | −0.41 | −0.80 | −1.29 | −1.95 | −0.25 | −0.39 | −0.49 | −0.66 |
| CNP | −0.06 | 0.07 | 0.37 | 1.31 | 1.94 | 0.13 | 0.30 | 0.93 | 0.63 |
| WIF1 | 0.01 | −0.24 | −0.51 | −0.94 | −1.74 | −0.25 | −0.27 | −0.43 | −0.80 |
| NPTX2 | −0.07 | −0.13 | −0.35 | −1.16 | −1.97 | −0.05 | −0.22 | −0.81 | −0.82 |
| LRFN2 | −0.04 | −0.16 | −0.24 | −0.92 | −1.76 | −0.12 | −0.08 | −0.68 | −0.83 |
| OMP | 0.31 | 0.33 | 0.69 | 0.68 | 1.00 | 0.02 | 0.36 | −0.01 | 0.32 |
| PCP4L1 | 0.14 | −0.19 | −0.78 | −1.05 | −1.49 | −0.33 | −0.58 | −0.28 | −0.43 |
| WIF1.1 | −0.01 | −0.21 | −0.48 | −0.91 | −1.69 | −0.21 | −0.27 | −0.43 | −0.78 |
| LRTM2 | −0.06 | −0.13 | −0.42 | −1.10 | −1.93 | −0.07 | −0.28 | −0.68 | −0.83 |
| ZFYVE27 | −0.10 | 0.08 | 0.00 | 1.42 | 0.06 | 0.18 | −0.07 | 1.42 | −1.36 |
| CPLX2 | −0.11 | −0.05 | −0.34 | −0.84 | −1.63 | 0.06 | −0.29 | −0.50 | −0.79 |
| DCC | −0.17 | −0.15 | −0.45 | −1.04 | −1.66 | 0.02 | −0.30 | −0.59 | −0.62 |
| TNFRSF8 | −0.17 | −0.34 | −0.81 | −0.52 | −0.67 | −0.16 | −0.47 | 0.29 | −0.15 |
| ATP5PF | 0.04 | 0.28 | 0.77 | 0.79 | 0.91 | 0.23 | 0.50 | 0.02 | 0.11 |
| WFDC2 | −0.08 | −0.17 | −0.66 | −1.23 | −1.95 | −0.09 | −0.49 | −0.58 | −0.71 |
| NPTXR | −0.04 | −0.10 | −0.22 | −0.99 | −1.86 | −0.07 | −0.12 | −0.77 | −0.87 |

**C** (Serum, Olink)

| | vs. HC: Early preHD | vs. HC: Late preHD | vs. HC: Early HD | vs. HC: Mod. HD | vs. HC: Adv. HD | Late/ Early preHD | Early HD/ Late preHD | Mod./ Early HD | Adv./ Mod. HD |
|---|---|---|---|---|---|---|---|---|---|
| NEFL | 0.32 | 1.00 | 1.58 | 1.81 | 2.36 | 0.68 | 0.58 | 0.23 | 0.55 |
| PRL | −0.01 | −0.06 | 0.27 | 1.20 | 2.79 | −0.04 | 0.33 | 0.92 | 1.60 |
| CD302 | 0.00 | 0.02 | 0.04 | 0.26 | 0.39 | 0.01 | 0.03 | 0.22 | 0.13 |
| GFRA1 | 0.01 | −0.05 | 0.02 | 0.32 | 0.40 | −0.07 | 0.07 | 0.30 | 0.08 |
| RNASE6 | −0.01 | 0.01 | 0.05 | 0.20 | 0.67 | 0.02 | 0.04 | 0.15 | 0.47 |
| TNFRSF19 | 0.10 | 0.03 | 0.10 | 0.23 | 0.46 | −0.06 | 0.06 | 0.13 | 0.23 |
| CHGA | 0.18 | 0.04 | 0.15 | 0.33 | 1.15 | −0.14 | 0.11 | 0.18 | 0.82 |
| XG | −0.01 | −0.07 | 0.00 | 0.20 | 0.44 | −0.07 | 0.07 | 0.20 | 0.24 |
| FBLN2 | 0.04 | −0.05 | 0.08 | 0.27 | 0.42 | −0.09 | 0.13 | 0.19 | 0.16 |
| CCER2 | 0.18 | 0.18 | 0.24 | 0.85 | 0.49 | −0.00 | 0.06 | 0.61 | −0.37 |
| DTNB | 0.05 | −0.13 | 0.08 | 0.24 | 0.58 | −0.18 | 0.21 | 0.16 | 0.34 |
| SFRP1 | −0.12 | −0.17 | −0.06 | 0.05 | 0.67 | −0.05 | 0.11 | 0.10 | 0.63 |
| FGF23 | −0.02 | −0.26 | 0.07 | 0.31 | 0.65 | −0.24 | 0.33 | 0.24 | 0.34 |
| PTGDS | −0.00 | −0.02 | 0.02 | 0.23 | 0.40 | −0.02 | 0.04 | 0.21 | 0.17 |
| F2R | 0.06 | −0.01 | 0.08 | 0.18 | 0.57 | −0.07 | 0.09 | 0.09 | 0.39 |
| RBFOX3 | 0.04 | −0.07 | 0.11 | 0.33 | 0.66 | −0.11 | 0.18 | 0.21 | 0.33 |
| GFAP | 0.09 | −0.03 | 0.15 | 0.43 | 0.74 | −0.12 | 0.18 | 0.28 | 0.32 |
| WFDC2 | 0.15 | 0.05 | 0.13 | 0.31 | 0.51 | −0.10 | 0.07 | 0.18 | 0.21 |
| CD276 | −0.01 | −0.01 | −0.00 | 0.26 | 0.55 | 0.00 | 0.01 | 0.26 | 0.30 |
| CHGB | 0.06 | −0.10 | 0.01 | 0.15 | 0.54 | −0.16 | 0.11 | 0.13 | 0.39 |
| LRG1 | −0.21 | −0.06 | 0.11 | 0.11 | 0.47 | 0.16 | 0.17 | −0.01 | 0.37 |
| OMG | −0.13 | −0.54 | −0.55 | −0.79 | −0.13 | −0.41 | −0.01 | −0.23 | 0.66 |
| IL17C | 0.04 | −0.15 | −0.02 | 0.19 | 1.02 | −0.19 | 0.13 | 0.21 | 0.83 |
| AGRP | 0.14 | −0.02 | 0.12 | 0.17 | 0.71 | −0.16 | 0.14 | 0.05 | 0.54 |
| PPL | −0.03 | −0.07 | −0.01 | 0.36 | 0.50 | −0.04 | 0.06 | 0.37 | 0.14 |

**D** (Serum, Soma)

| | vs. HC: Early preHD | vs. HC: Late preHD | vs. HC: Early HD | vs. HC: Mod. HD | vs. HC: Adv. HD | Late/ Early preHD | Early HD/ Late preHD | Mod./ Early HD | Adv./ Mod. HD |
|---|---|---|---|---|---|---|---|---|---|
| PRL | 0.06 | −0.09 | 0.19 | 1.19 | 2.70 | −0.15 | 0.28 | 1.00 | 1.50 |
| PRL.1 | 0.01 | −0.08 | 0.22 | 1.07 | 2.43 | −0.09 | 0.30 | 0.85 | 1.36 |
| PATE1 | 0.15 | −0.08 | −0.02 | 0.07 | 2.14 | −0.24 | 0.06 | 0.09 | 2.07 |
| CARD17 | −0.19 | 0.12 | 0.12 | 0.15 | 1.97 | 0.31 | 0.01 | 0.03 | 1.82 |
| CRLF2 | 0.04 | 0.01 | 0.18 | 0.24 | 1.90 | −0.03 | 0.17 | 0.06 | 1.65 |
| CRYGC | −0.18 | −0.02 | 0.12 | 0.15 | 1.93 | 0.16 | 0.13 | 0.04 | 1.77 |
| ACP3 | −0.01 | −0.15 | 0.04 | 1.45 | 0.07 | −0.14 | 0.19 | 1.42 | −1.38 |
| TMPRSS15 | −0.14 | −0.15 | −0.11 | −0.06 | 1.62 | −0.01 | 0.05 | 0.05 | 1.68 |
| KRAS | 0.03 | −0.08 | 0.06 | 0.02 | 1.85 | −0.11 | 0.14 | −0.03 | 1.82 |
| AARSD1 | −0.13 | 0.18 | 0.17 | 0.31 | 1.60 | 0.30 | −0.01 | 0.15 | 1.29 |
| IL10RB | −0.17 | 0.11 | 0.23 | 0.62 | 1.57 | 0.28 | 0.12 | 0.39 | 0.95 |
| FGF8 | −0.20 | 0.15 | 0.15 | 0.45 | 1.55 | 0.35 | −0.00 | 0.31 | 1.10 |
| SPAG11A | 0.01 | 0.15 | 0.17 | 0.16 | 1.41 | 0.14 | 0.03 | −0.01 | 1.25 |
| CALCB | −0.01 | −0.08 | −0.26 | −0.19 | 1.30 | −0.08 | −0.18 | 0.07 | 1.49 |
| ANTXR2 | 0.09 | 0.04 | 0.18 | −0.58 | −1.40 | −0.05 | 0.14 | −0.76 | −0.82 |
| KCTD15 | −0.11 | −0.00 | 0.07 | 0.12 | 1.50 | 0.11 | 0.07 | 0.05 | 1.38 |
| UBE2S | 0.06 | −0.43 | −0.04 | −0.20 | 1.27 | −0.49 | 0.39 | −0.17 | 1.47 |
| TNFRSF10D | −0.01 | 0.20 | 0.05 | 0.37 | 1.72 | 0.21 | −0.15 | 0.33 | 1.35 |
| ENTPD3 | −0.37 | 0.01 | 0.15 | 0.40 | 1.51 | 0.38 | 0.13 | 0.26 | 1.11 |
| TXLNA | −0.04 | 0.14 | 0.10 | 0.09 | 1.46 | 0.18 | −0.04 | −0.00 | 1.37 |
| MLX | −0.16 | 0.06 | 0.13 | 0.19 | 1.39 | 0.22 | 0.07 | 0.07 | 1.20 |
| SAA2 | −0.11 | 0.05 | −0.05 | 0.18 | 1.47 | 0.16 | −0.09 | 0.23 | 1.29 |
| HEY1 | 0.04 | 0.20 | 0.19 | 0.21 | 1.61 | 0.15 | −0.01 | 0.02 | 1.40 |
| CLEC7A | −0.02 | 0.13 | 0.07 | 0.02 | 1.54 | 0.15 | −0.06 | −0.05 | 1.52 |
| AHSA1 | −0.28 | −0.01 | 0.10 | 0.29 | 1.36 | 0.27 | 0.11 | 0.19 | 1.07 |

FDR: >0.05, <0.05, 0.01, 0.001, 0.0001, 0

**Fig. S3. Differential abundance of CSF and serum proteins across HD-ISS stages.** (**A**-**D**) Heatmaps showing the top 25 differentially abundant proteins from (A) Olink CSF, (B) Soma CSF, (C) Olink serum, and (D) Soma serum across pairwise HD-ISS stage contrasts. Multiple linear regression models were adjusted for age and sex. HD-ISS stages for all HDGECs were imputed as in *(31)* based on landmarks defined in *(30)*. Columns represent group contrasts, and rows show proteins ordered first by the number of significant contrasts (FDR-adjusted $p < 0.05$) and then by minimum adjusted *p*-value. Cell values correspond to $\log_2$ fold changes, with text indicating the direction and magnitude of change, while colour intensity reflects statistical significance (FDR-adjusted *p*-value). Shared CSF proteins between Olink and Soma platforms are shown in blue, and shared serum proteins are shown in red. Shared proteins between biofluid compartments measured on the same platform are shown in bold/purple.

**Olink** / **Soma**

**A** (CSF, Olink)

| | vs. HC | | | | Stage 1 | Stage 2 | Stage 3 |
|---|---|---|---|---|---|---|---|
| | Stage 0 | Stage 1 | Stage 2 | Stage 3 | vs. 0 | vs. 1 | vs. 2 |
| NEFL | 1.57 | 2.25 | 2.08 | 2.58 | 0.69 | −0.17 | 0.50 |
| TNFRSF8 | −0.42 | −0.72 | −1.10 | −1.70 | −0.30 | −0.38 | −0.60 |
| TPM3 | 0.27 | 0.47 | 0.48 | 0.76 | 0.20 | 0.01 | 0.28 |
| NPPB | −0.10 | −0.14 | −0.36 | −0.55 | −0.04 | −0.22 | −0.19 |
| WFDC2 | −0.03 | −0.10 | −0.11 | −0.24 | −0.07 | −0.01 | −0.13 |
| SCN4B | −0.06 | −0.05 | −0.18 | −0.34 | 0.01 | −0.13 | −0.16 |
| MME | −0.05 | −0.07 | −0.22 | −0.36 | −0.02 | −0.15 | −0.14 |
| DNM3 | 0.03 | 0.15 | 0.22 | 0.24 | 0.11 | 0.07 | 0.02 |
| CHI3L1 | 0.18 | 0.30 | 0.23 | 0.29 | 0.11 | −0.06 | 0.05 |
| OMP | 0.34 | 0.14 | 0.22 | 0.47 | −0.20 | 0.08 | 0.25 |
| PRL | 0.23 | 0.15 | 0.06 | 0.52 | −0.08 | −0.09 | 0.46 |
| GPR101 | −0.04 | −0.21 | −0.29 | −0.23 | −0.17 | −0.08 | 0.06 |
| ATP5IF1 | 0.16 | 0.15 | 0.25 | 0.25 | −0.01 | 0.10 | 0.01 |
| LRTM2 | −0.04 | −0.10 | −0.14 | −0.38 | −0.06 | −0.04 | −0.24 |
| WIF1 | −0.10 | −0.07 | −0.12 | −0.42 | 0.03 | −0.05 | −0.30 |
| TNN | −0.14 | −0.01 | −0.34 | −0.42 | 0.13 | −0.33 | −0.08 |
| NDUFS6 | 0.08 | 0.22 | 0.06 | 0.28 | 0.15 | −0.17 | 0.22 |
| MFGE8 | −0.17 | −0.10 | −0.10 | −0.26 | 0.08 | −0.00 | −0.16 |
| TSC1 | −0.13 | −0.03 | −0.59 | −0.14 | 0.10 | −0.55 | 0.44 |
| SERPINA3 | 0.04 | 0.23 | 0.11 | 0.14 | 0.19 | −0.12 | 0.03 |
| SMPD3 | −0.09 | −0.21 | −0.12 | −0.21 | −0.12 | 0.09 | −0.08 |
| COCH | −0.00 | 0.03 | −0.05 | −0.08 | 0.03 | −0.07 | −0.04 |
| NPTX2 | −0.00 | −0.03 | −0.15 | −0.26 | −0.03 | −0.12 | −0.12 |
| PLA2G7 | 0.09 | −0.13 | −0.09 | −0.20 | −0.22 | 0.04 | −0.11 |
| CHIT1 | 1.00 | 1.36 | 0.26 | 0.76 | 0.36 | −1.11 | 0.51 |

**B** (CSF, Soma)

| | vs. HC | | | | Stage 1 | Stage 2 | Stage 3 |
|---|---|---|---|---|---|---|---|
| | Stage 0 | Stage 1 | Stage 2 | Stage 3 | vs. 0 | vs. 1 | vs. 2 |
| NEFL | 0.90 | 1.48 | 1.49 | 1.89 | 0.58 | 0.01 | 0.40 |
| NEFH | 0.71 | 1.24 | 1.20 | 1.64 | 0.53 | −0.04 | 0.44 |
| ITGA1/ITGB1 | 0.52 | 0.58 | 0.97 | 1.01 | 0.06 | 0.40 | 0.04 |
| SEMA3E | −0.04 | −0.17 | −1.05 | −0.21 | −0.13 | −0.88 | 0.84 |
| NPPB | −0.23 | −0.34 | −0.62 | −1.10 | −0.11 | −0.28 | −0.48 |
| MAP4K1 | 0.30 | 0.71 | 0.70 | 1.08 | 0.40 | −0.01 | 0.39 |
| PDE1B | −0.17 | −0.46 | −0.52 | −1.04 | −0.28 | −0.06 | −0.51 |
| GFAP | 0.21 | 0.55 | 0.56 | 0.78 | 0.34 | 0.00 | 0.23 |
| PCP4L1 | 0.16 | −0.26 | −0.30 | −0.90 | −0.42 | −0.04 | −0.60 |
| LINGO3 | −0.36 | −0.34 | −0.58 | −0.87 | 0.02 | −0.24 | −0.29 |
| WFDC2 | 0.09 | −0.41 | −0.32 | −0.79 | −0.50 | 0.09 | −0.47 |
| ATP5PF | 0.08 | 0.28 | 0.35 | 0.78 | 0.19 | 0.07 | 0.43 |
| TNFRSF8 | −0.14 | −0.30 | −0.60 | −0.76 | −0.16 | −0.30 | −0.16 |
| MAPRE3 | 0.34 | 0.46 | 0.63 | 0.63 | 0.12 | 0.18 | −0.00 |
| OMP | 0.46 | 0.27 | 0.40 | 0.65 | −0.19 | 0.12 | 0.26 |
| COX5A | 0.29 | 0.40 | 0.56 | 0.61 | 0.10 | 0.17 | 0.05 |
| WIF1 | −0.13 | −0.22 | −0.15 | −0.62 | −0.09 | 0.07 | −0.47 |
| PRL | 0.18 | 0.06 | 0.09 | 0.61 | −0.11 | 0.03 | 0.52 |
| WIF1.1 | −0.12 | −0.20 | −0.12 | −0.59 | −0.08 | 0.07 | −0.47 |
| PTPA | 0.39 | 0.51 | 0.59 | 0.53 | 0.12 | 0.09 | −0.07 |
| TESC | −0.19 | −0.33 | −0.28 | −0.62 | −0.14 | 0.05 | −0.34 |
| WNT16 | −0.30 | −0.18 | −0.19 | −0.60 | 0.13 | −0.02 | −0.41 |
| DCC | −0.13 | −0.19 | −0.26 | −0.57 | −0.06 | −0.07 | −0.31 |
| LRTM2 | −0.03 | −0.23 | −0.20 | −0.56 | −0.20 | 0.03 | −0.36 |
| GARS1 | 0.35 | 0.38 | 0.56 | 0.49 | 0.04 | 0.18 | −0.07 |

**C** (Serum, Olink)

| | vs. HC | | | | Stage 1 | Stage 2 | Stage 3 |
|---|---|---|---|---|---|---|---|
| | Stage 0 | Stage 1 | Stage 2 | Stage 3 | vs. 0 | vs. 1 | vs. 2 |
| **NEFL** | 0.58 | 1.03 | 1.01 | 1.63 | 0.45 | −0.02 | 0.62 |
| AHNAK | −0.06 | −0.08 | 0.07 | 0.22 | −0.02 | 0.15 | 0.15 |
| OMG | −0.43 | −0.33 | −0.49 | −0.54 | 0.11 | −0.16 | −0.06 |
| NCAN | 0.00 | −0.21 | −0.22 | −0.21 | −0.22 | −0.01 | 0.01 |
| SCRG1 | 0.06 | 0.00 | 0.03 | 0.18 | −0.05 | 0.02 | 0.15 |
| ACP5 | 0.08 | 0.05 | 0.15 | 0.19 | −0.04 | 0.11 | 0.04 |
| TRIM21 | 0.06 | 0.14 | 0.14 | 0.28 | 0.08 | 0.00 | 0.14 |
| SPRR3 | 0.09 | 0.13 | 0.26 | 0.34 | 0.04 | 0.13 | 0.09 |
| CYTL1 | 0.08 | −0.00 | 0.11 | 0.17 | −0.09 | 0.11 | 0.07 |
| APLP1 | −0.06 | −0.15 | −0.12 | −0.23 | −0.09 | 0.03 | −0.10 |
| EFNA4 | 0.07 | −0.05 | 0.15 | 0.14 | −0.11 | 0.20 | −0.01 |
| CCER2 | 0.18 | −0.06 | 0.27 | 0.36 | −0.24 | 0.33 | 0.08 |
| OSCAR | 0.09 | −0.04 | 0.17 | 0.17 | −0.13 | 0.21 | −0.00 |
| OGFR | 0.03 | −0.02 | 0.07 | 0.16 | −0.05 | 0.09 | 0.09 |
| TPPP3 | 0.03 | 0.03 | 0.20 | 0.37 | 0.00 | 0.16 | 0.17 |
| LGALS3 | 0.03 | −0.00 | 0.12 | 0.17 | −0.03 | 0.12 | 0.04 |
| ADAMTS8 | −0.06 | −0.12 | −0.08 | −0.20 | −0.06 | 0.04 | −0.12 |
| KRT19 | −0.09 | −0.09 | 0.28 | 0.26 | 0.01 | 0.37 | −0.02 |
| DSCAM | 0.19 | 0.07 | 0.24 | 0.23 | −0.12 | 0.18 | −0.02 |
| IL15RA | 0.06 | 0.03 | 0.07 | 0.15 | −0.04 | 0.05 | 0.08 |
| VWC2 | 0.03 | −0.05 | 0.08 | 0.16 | −0.08 | 0.13 | 0.09 |
| LILRB5 | −0.57 | −0.34 | −0.75 | −0.20 | 0.23 | −0.40 | 0.55 |
| ANXA10 | −0.23 | −0.29 | 1.00 | 0.55 | −0.06 | 1.29 | −0.45 |
| MST1 | −0.06 | 0.03 | −0.52 | 0.03 | 0.09 | −0.55 | 0.56 |
| SLC9A3R2 | −0.03 | −0.37 | 0.10 | 0.01 | −0.34 | 0.47 | −0.09 |

**D** (Serum, Soma)

| | vs. HC | | | | Stage 1 | Stage 2 | Stage 3 |
|---|---|---|---|---|---|---|---|
| | Stage 0 | Stage 1 | Stage 2 | Stage 3 | vs. 0 | vs. 1 | vs. 2 |
| OMG | −0.58 | −0.36 | −0.67 | −0.69 | 0.22 | −0.31 | −0.02 |
| TMEM132B | −0.31 | −0.35 | −0.24 | −0.58 | −0.03 | 0.11 | −0.34 |
| NCAN | 0.05 | −0.36 | −0.66 | −0.57 | −0.41 | −0.30 | 0.09 |
| NPTXR | −0.35 | −0.24 | −0.53 | −0.59 | 0.11 | −0.29 | −0.06 |
| NPTXR.1 | −0.33 | −0.41 | −0.55 | −0.57 | −0.07 | −0.14 | −0.02 |
| MAG | −0.24 | −0.35 | −0.45 | −0.55 | −0.11 | −0.10 | −0.10 |
| FXYD2 | 0.10 | 0.22 | 0.31 | 0.52 | 0.12 | 0.10 | 0.21 |
| NRCAM | −0.23 | −0.15 | −0.19 | −0.47 | 0.08 | −0.04 | −0.28 |
| ADAM23 | −0.13 | −0.26 | −0.39 | −0.49 | −0.13 | −0.13 | −0.10 |
| GOT2 | −0.47 | −0.33 | −0.45 | −0.49 | 0.14 | −0.12 | −0.04 |
| SORCS1 | −0.33 | −0.34 | −0.49 | −0.47 | −0.01 | −0.15 | 0.03 |
| USF2 | 0.63 | 0.03 | 0.01 | 0.02 | −0.60 | −0.02 | 0.00 |
| KIN | 0.73 | 0.05 | −0.03 | −0.01 | −0.68 | −0.08 | 0.02 |
| TMEM225B | 0.04 | 0.72 | 0.07 | 0.12 | 0.68 | −0.64 | 0.05 |
| SHANK3 | −0.34 | −0.77 | −0.12 | −0.32 | −0.42 | 0.64 | −0.20 |
| FAM204A | 0.15 | 0.71 | 0.07 | 0.05 | 0.57 | −0.65 | −0.02 |
| GSTZ1 | −0.54 | −0.79 | −0.62 | −0.28 | −0.25 | 0.16 | 0.34 |
| SETMAR | 0.00 | 0.78 | 0.23 | −0.09 | 0.78 | −0.56 | −0.32 |
| SORT1 | 0.29 | 0.09 | 0.65 | 0.12 | −0.20 | 0.56 | −0.53 |
| DEFB125 | 0.21 | 0.00 | 0.86 | 0.12 | −0.21 | 0.86 | −0.74 |
| LILRB5 | −0.59 | −0.27 | −0.76 | −0.26 | 0.32 | −0.49 | 0.50 |
| KLKB1 | −0.14 | −0.02 | −0.74 | −0.16 | 0.12 | −0.72 | 0.58 |
| UBE2T | 0.14 | 0.14 | 0.55 | 0.08 | −0.00 | 0.42 | −0.47 |
| MANBA | 0.00 | −0.21 | 0.47 | −0.22 | −0.21 | 0.68 | −0.69 |
| LECT2 | −0.23 | −0.25 | −0.72 | 0.02 | −0.02 | −0.47 | 0.74 |

**FDR-adjusted *p*-value**
—>0.05
<0.05
0.01
0.001
0.0001
0

**Fig. S4. CSF-serum protein correlations across analytical platforms.** (**A-D**) Balloon plots illustrating partial correlations between CSF and serum protein levels, adjusted for age and sex. The x-axis shows the partial Pearson's correlation coefficient (r), and the y-axis lists the corresponding proteins. Balloon size and colour intensity reflect the statistical significance of the correlation, represented as $-\log_{10}$ FDR-adjusted *p*-values. Top 25 proteins with the strongest CSF–serum correlations identified using the (A) Olink and (B) Soma platforms, respectively. Top 25 differentially abundant proteins in both CSF and serum with concordant directional changes identified using (C) Olink and (D) Soma platforms, respectively. Proteins were independently ranked by significance (FDR-adjusted *p*-value) within CSF and serum and plotted based on their rank difference between biofluids to emphasize biofluid-specific and concordant expression patterns. Differentially abundant proteins in both CSF and serum shared across platforms are bolded.

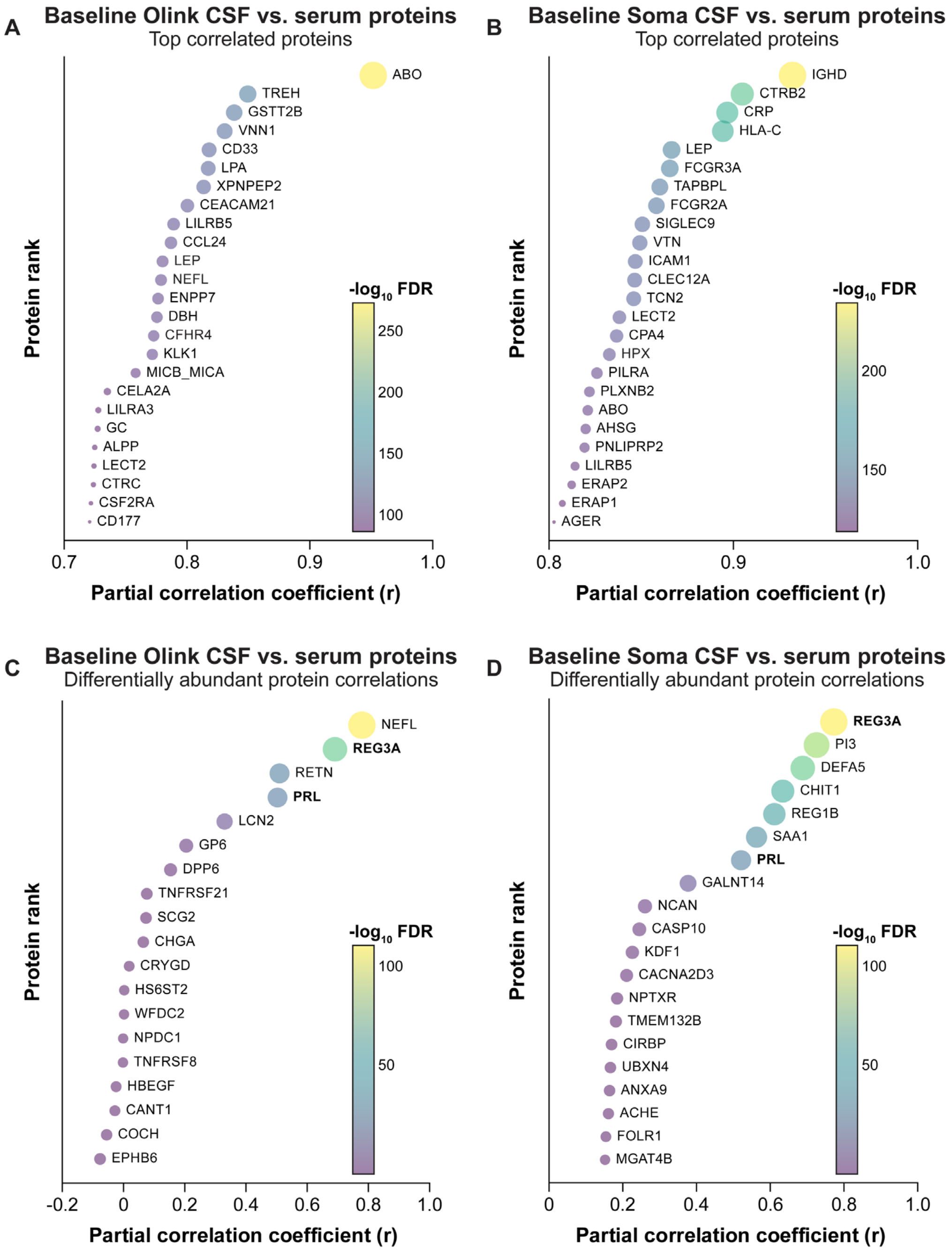

**Fig. S5. Associations of baseline CSF and serum protein abundance with clinical measures of HD.** (A-D) Heatmaps showing the top 25 correlations between protein abundance from (A) Olink CSF, (B) Soma CSF, (C) Olink serum, and (D) Soma serum, and clinical measures of disease severity in HDGEC. Partial Spearman models were adjusted for age and sex. Columns represent clinical measures and rows display proteins ordered first by the number of significant correlations (FDR < 0.05) and then by minimum FDR. Cell values correspond to partial Spearman correlation coefficients (ρ), with text indicating the strength and direction of association, while colour intensity reflects statistical significance (FDR). Shared CSF proteins between Olink and Soma platforms are shown in blue, and shared serum proteins are shown in red. Shared proteins between biofluid compartments measured on the same platform are shown in bold/purple.

**Olink** | **Soma**

FDR: >0.05; <0.05; 0.01; 0.001; 0.0001; 0

**A** (Olink, CSF)

| | SCNT | SDMT | SWRT | TFC | TMS | VFT | cUHDRS |
|---|---|---|---|---|---|---|---|
| TNFRSF8 | 0.54 | 0.55 | 0.53 | 0.43 | −0.66 | 0.46 | 0.60 |
| NEFL | −0.38 | −0.38 | −0.35 | −0.31 | 0.46 | −0.36 | −0.43 |
| NPPB | 0.38 | 0.30 | 0.39 | 0.31 | −0.44 | 0.24 | 0.39 |
| TPM3 | −0.35 | −0.36 | −0.34 | −0.34 | 0.43 | −0.33 | −0.41 |
| WFDC2 | 0.29 | 0.26 | 0.27 | 0.24 | −0.33 | 0.20 | 0.29 |
| LPO | 0.31 | 0.19 | 0.30 | 0.18 | −0.30 | 0.13 | 0.28 |
| COCH | 0.24 | 0.19 | 0.25 | 0.21 | −0.28 | 0.14 | 0.25 |
| MME | 0.27 | 0.22 | 0.24 | 0.16 | −0.27 | 0.18 | 0.24 |
| LRTM2 | 0.24 | 0.17 | 0.22 | 0.20 | −0.26 | 0.14 | 0.22 |
| SCN4B | 0.21 | 0.19 | 0.19 | 0.18 | −0.26 | 0.15 | 0.23 |
| CHGA | 0.26 | 0.18 | 0.24 | 0.18 | −0.23 | 0.11 | 0.22 |
| WIF1 | 0.23 | 0.17 | 0.24 | 0.18 | −0.25 | 0.14 | 0.22 |
| MMP7 | −0.20 | −0.17 | −0.15 | −0.16 | 0.20 | −0.25 | −0.20 |
| SPON2 | 0.20 | 0.15 | 0.23 | 0.13 | −0.23 | 0.10 | 0.21 |
| ADAM23 | 0.21 | 0.14 | 0.21 | 0.16 | −0.23 | 0.11 | 0.21 |
| NCR3LG1 | 0.18 | 0.12 | 0.18 | 0.15 | −0.22 | 0.09 | 0.18 |
| DNM3 | −0.15 | −0.21 | −0.14 | −0.14 | 0.22 | −0.17 | −0.21 |
| PRL | −0.14 | −0.12 | −0.11 | −0.21 | 0.18 | −0.17 | −0.18 |
| ADAMTS4 | 0.18 | 0.14 | 0.21 | 0.14 | −0.19 | 0.12 | 0.18 |
| ASAH2 | 0.17 | 0.14 | 0.21 | 0.10 | −0.19 | 0.04 | 0.19 |
| TARBP2 | 0.21 | 0.17 | 0.17 | 0.16 | −0.10 | 0.16 | 0.18 |
| NPTX2 | 0.21 | 0.16 | 0.20 | 0.16 | −0.21 | 0.09 | 0.19 |
| PDAP1 | 0.21 | 0.18 | 0.21 | 0.12 | −0.17 | 0.13 | 0.19 |
| TCTN3 | 0.20 | 0.11 | 0.20 | 0.12 | −0.21 | 0.06 | 0.16 |
| ECM1 | 0.21 | 0.11 | 0.20 | 0.11 | −0.19 | 0.14 | 0.16 |

**B** (Soma, CSF)

| | SCNT | SDMT | SWRT | TFC | TMS | VFT | cUHDRS |
|---|---|---|---|---|---|---|---|
| NEFL | −0.51 | −0.48 | −0.47 | −0.39 | 0.57 | −0.43 | −0.55 |
| NEFH | −0.48 | −0.44 | −0.44 | −0.37 | 0.52 | −0.42 | −0.51 |
| NPPB | 0.36 | 0.30 | 0.39 | 0.36 | −0.46 | 0.26 | 0.42 |
| MAP4K1 | −0.42 | −0.36 | −0.42 | −0.27 | 0.42 | −0.29 | −0.43 |
| PDE1B | 0.32 | 0.30 | 0.33 | 0.26 | −0.41 | 0.23 | 0.35 |
| WFDC2 | 0.33 | 0.34 | 0.30 | 0.28 | −0.38 | 0.26 | 0.35 |
| PCP4L1 | 0.30 | 0.31 | 0.27 | 0.29 | −0.35 | 0.21 | 0.33 |
| CNP | −0.28 | −0.25 | −0.30 | −0.29 | 0.31 | −0.22 | −0.34 |
| GFAP | −0.27 | −0.27 | −0.22 | −0.21 | 0.31 | −0.26 | −0.30 |
| LINGO3 | 0.22 | 0.20 | 0.23 | 0.22 | −0.30 | 0.18 | 0.27 |
| WIF1 | 0.26 | 0.21 | 0.27 | 0.23 | −0.28 | 0.19 | 0.26 |
| ATP5PF | −0.20 | −0.21 | −0.14 | −0.21 | 0.28 | −0.11 | −0.24 |
| WIF1.1 | 0.26 | 0.21 | 0.27 | 0.22 | −0.28 | 0.18 | 0.26 |
| MMP7 | −0.25 | −0.22 | −0.19 | −0.20 | 0.26 | −0.24 | −0.24 |
| TNFRSF8 | 0.18 | 0.24 | 0.25 | 0.11 | −0.26 | 0.10 | 0.25 |
| NPTX2 | 0.25 | 0.20 | 0.23 | 0.23 | −0.25 | 0.16 | 0.24 |
| MRPL12 | −0.21 | −0.20 | −0.19 | −0.24 | 0.24 | −0.06 | −0.25 |
| PPP1R1A | 0.22 | 0.22 | 0.22 | 0.17 | −0.24 | 0.07 | 0.23 |
| ITPA | 0.21 | 0.18 | 0.24 | 0.19 | −0.24 | 0.17 | 0.23 |
| B3GALT2 | 0.20 | 0.18 | 0.18 | 0.22 | −0.24 | 0.12 | 0.22 |
| TCEAL3 | −0.23 | −0.18 | −0.24 | −0.11 | 0.23 | −0.14 | −0.23 |
| MDK | 0.22 | 0.22 | 0.19 | 0.17 | −0.22 | 0.16 | 0.24 |
| B3GNT2 | 0.21 | 0.14 | 0.22 | 0.16 | −0.23 | 0.10 | 0.20 |
| TESC | 0.20 | 0.19 | 0.23 | 0.21 | −0.23 | 0.15 | 0.23 |
| TNFRSF18 | 0.21 | 0.17 | 0.22 | 0.13 | −0.22 | 0.15 | 0.21 |

**C** (Olink, Serum)

| | SCNT | SDMT | SWRT | TFC | TMS | VFT | cUHDRS |
|---|---|---|---|---|---|---|---|
| **NEFL** | −0.46 | −0.48 | −0.42 | −0.42 | 0.58 | −0.40 | −0.53 |
| PALM | −0.25 | −0.17 | −0.24 | −0.32 | 0.29 | −0.23 | −0.28 |
| PHLDB1 | −0.20 | −0.16 | −0.19 | −0.30 | 0.28 | −0.17 | −0.25 |
| DMD | −0.20 | −0.17 | −0.16 | −0.29 | 0.26 | −0.14 | −0.24 |
| FBLN2 | −0.22 | −0.13 | −0.21 | −0.29 | 0.21 | −0.15 | −0.23 |
| ENAH | −0.25 | −0.18 | −0.22 | −0.25 | 0.28 | −0.22 | −0.26 |
| SCARA5 | −0.26 | −0.21 | −0.27 | −0.27 | 0.27 | −0.19 | −0.28 |
| AHNAK2 | −0.22 | −0.13 | −0.19 | −0.23 | 0.28 | −0.19 | −0.23 |
| PALM2 | −0.22 | −0.14 | −0.20 | −0.27 | 0.21 | −0.20 | −0.23 |
| ELN | −0.22 | −0.21 | −0.21 | −0.22 | 0.27 | −0.21 | −0.25 |
| NUCB2 | −0.18 | −0.12 | −0.19 | −0.26 | 0.25 | −0.13 | −0.22 |
| AHNAK | −0.21 | −0.16 | −0.22 | −0.25 | 0.25 | −0.12 | −0.25 |
| GFRA1 | −0.20 | −0.13 | −0.19 | −0.25 | 0.14 | −0.12 | −0.21 |
| DPY30 | −0.21 | −0.15 | −0.22 | −0.24 | 0.18 | −0.12 | −0.23 |
| A1BG | −0.20 | −0.13 | −0.17 | −0.19 | 0.23 | −0.17 | −0.19 |
| CRNN | −0.20 | −0.12 | −0.19 | −0.20 | 0.23 | −0.09 | −0.20 |
| GGH | −0.23 | −0.11 | −0.23 | −0.19 | 0.17 | −0.11 | −0.20 |
| EFHD1 | −0.20 | −0.15 | −0.14 | −0.23 | 0.22 | −0.16 | −0.20 |
| LRG1 | −0.20 | −0.19 | −0.17 | −0.22 | 0.20 | −0.18 | −0.23 |
| C9 | −0.23 | −0.20 | −0.21 | −0.15 | 0.18 | −0.10 | −0.21 |
| VASN | −0.21 | −0.19 | −0.15 | −0.20 | 0.23 | −0.17 | −0.22 |
| GFAP | −0.18 | −0.19 | −0.19 | −0.18 | 0.17 | −0.18 | −0.21 |
| CKAP4 | −0.20 | −0.13 | −0.21 | −0.19 | 0.17 | −0.14 | −0.19 |
| VWC2 | −0.20 | −0.20 | −0.19 | −0.21 | 0.14 | −0.16 | −0.19 |
| GPHA2 | −0.11 | −0.18 | −0.12 | −0.19 | 0.20 | −0.16 | −0.19 |

**D** (Soma, Serum)

| | SCNT | SDMT | SWRT | TFC | TMS | VFT | cUHDRS |
|---|---|---|---|---|---|---|---|
| DSG1 | −0.18 | −0.18 | −0.16 | −0.26 | 0.19 | −0.14 | −0.24 |
| JAM2 | −0.14 | −0.19 | −0.18 | −0.26 | 0.17 | −0.16 | −0.22 |
| LAYN | −0.16 | −0.21 | −0.16 | −0.25 | 0.19 | −0.10 | −0.24 |
| AFM | 0.22 | 0.19 | 0.21 | 0.22 | −0.23 | 0.18 | 0.25 |
| RNF215 | 0.22 | 0.16 | 0.17 | 0.17 | −0.24 | 0.10 | 0.21 |
| ALB | 0.20 | 0.18 | 0.23 | 0.18 | −0.23 | 0.16 | 0.24 |
| AFM.1 | 0.21 | 0.19 | 0.20 | 0.19 | −0.20 | 0.17 | 0.24 |
| APLP1 | 0.14 | 0.17 | 0.16 | 0.19 | −0.24 | 0.14 | 0.23 |
| CTNNBIP1 | −0.15 | −0.14 | −0.14 | −0.23 | 0.20 | −0.15 | −0.21 |
| ACVR2A | −0.13 | −0.18 | −0.13 | −0.21 | 0.23 | −0.14 | −0.22 |
| NRCAM | 0.22 | 0.17 | 0.14 | 0.22 | −0.22 | 0.23 | 0.19 |
| DEFB1 | 0.22 | 0.18 | 0.21 | 0.18 | −0.14 | 0.14 | 0.19 |
| PRKCD | 0.16 | 0.20 | 0.18 | 0.20 | −0.20 | 0.09 | 0.22 |
| PTPRJ | 0.16 | 0.17 | 0.18 | 0.21 | −0.20 | 0.17 | 0.20 |
| ARF3 | 0.21 | 0.21 | 0.20 | 0.17 | −0.12 | 0.17 | 0.20 |
| ANTKMT | 0.21 | 0.21 | 0.20 | 0.15 | −0.15 | 0.12 | 0.20 |
| VASN | −0.17 | −0.18 | −0.13 | −0.21 | 0.19 | −0.15 | −0.20 |
| HMGB2 | 0.17 | 0.19 | 0.18 | 0.21 | −0.17 | 0.12 | 0.21 |
| MRFAP1L1 | 0.19 | 0.14 | 0.21 | 0.11 | −0.14 | 0.18 | 0.16 |
| NCAN | 0.15 | 0.20 | 0.12 | 0.14 | −0.20 | 0.14 | 0.20 |
| ZNF41 | 0.19 | 0.19 | 0.18 | 0.18 | −0.15 | 0.09 | 0.20 |
| C9 | −0.20 | −0.15 | −0.19 | −0.16 | 0.14 | −0.11 | −0.19 |
| GGH | −0.19 | −0.10 | −0.19 | −0.18 | 0.12 | −0.12 | −0.18 |
| NPPA | 0.19 | 0.13 | 0.19 | 0.19 | −0.15 | 0.18 | 0.19 |
| C9.1 | −0.19 | −0.15 | −0.19 | −0.16 | 0.15 | −0.10 | −0.19 |

**Fig. S6. Associations of baseline CSF and serum protein abundance with YTO.** (**A**, **B**) Volcano plots showing the correlations of (A) CSF and (B) serum proteins at baseline with estimated years to predicted disease onset (YTO) in preHD measured using Olink and Soma platforms. Partial Pearson correlation coefficients (r) are plotted against FDR-adjusted *p*-values (−$\log_{10}$ transformed). Olink proteins are depicted as triangles, whereas Soma proteins are depicted as squares. Proteins showing significant inverse correlations with YTO in each biofluid are shown in blue, while those proteins with significant positive correlations are shown in red (FDR < 0.01). The top most significantly correlated Olink (orange) and Soma (green) proteins are annotated. Horizontal dotted line indicates the significance threshold. (**C**-**F**) Sample-level heatmaps showing correlations of the top 25 proteins at baseline from (C) Olink CSF, (D) Soma CSF, (E) Olink serum, and (F) Soma serum with YTO in preHD. Linear models were adjusted for sex. Columns represent proteins ordered by significance (FDR-adjusted *p*-value), and rows correspond to individual samples ordered by YTO. Cell values indicate scaled protein abundance levels (z-scores), and annotations display YTO and corresponding FDR-adjusted *p*-values (FDR) for each protein. Shared CSF proteins between Olink and Soma platforms are shown in blue, and shared serum proteins are shown in red. Shared proteins between biofluid compartments measured on the same platform are shown in bold/purple.

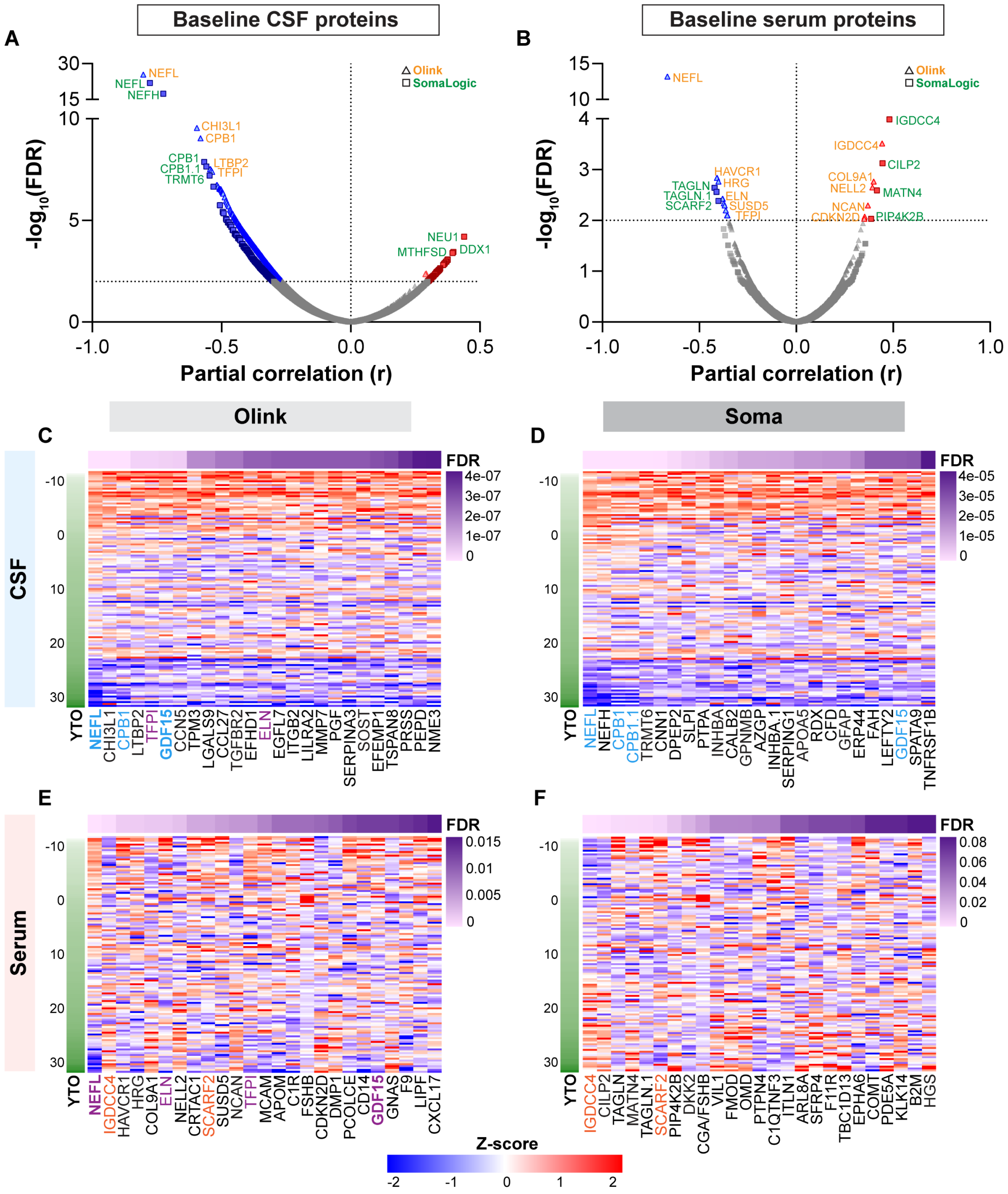

**Fig. S7. Functional enrichment analysis of CSF and serum proteins associated with YTO.** (**A-D**) Balloon plots illustrating enrichment of the top 10 GO Biological Process terms among proteins significantly associated with YTO in preHD for (A) Olink CSF, (B) Soma CSF, (C) Olink serum, and (D) Soma serum. The y-axis shows the top 10 enriched Biological Process terms, and x-axis shows -log10 FDR-adjusted *p*-value. Enrichment was performed using gProfiler. Proteins with nominal *p*-values <0.001 from partial correlation analysis (**Fig. S5**) were used as input. The term size cutoff was set to 2,500. Balloon colour indicates term size, and balloon size reflects intersection size.

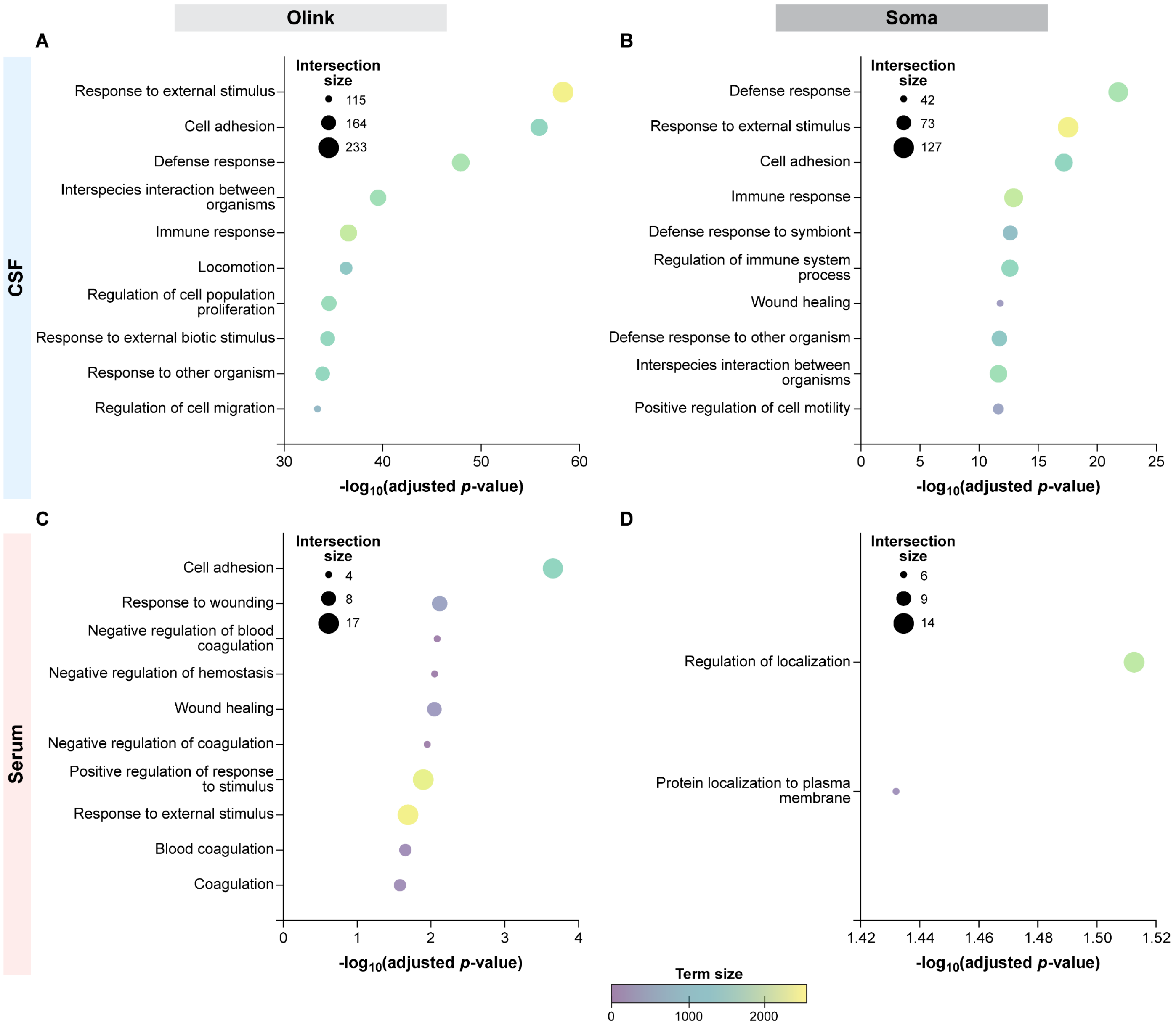
